# A water compartment cell culture lid enables stable longitudinal recording of neuronal networks in vitro

**DOI:** 10.64898/2026.04.01.713917

**Authors:** Benedikt Maurer, Fabio Fischer, Giulia Amos, Vaiva Vasiliauskaitė, János Vörös

**Affiliations:** Laboratory of Biosensors and Bioelectronics, Institute for Biomedical Engineering, University and ETH Zurich, Zurich, Switzerland

**Keywords:** neuronal networks, development, microelectrode arrays, *in vitro* neuro-science

## Abstract

Longitudinal electrophysiological recordings of neuronal networks are essential for studying network maturation, plasticity, and pharmacological responses. Yet current microelectrode array (MEA) approaches are limited by evaporation-induced drift in culture conditions, exacerbated by heat dissipation from active recording electronics on CMOS-based high-density MEAs. We present a cell culture lid featuring a water compartment at its interface that eliminates evaporation whilst maintaining gas exchange. Combined with a custom incubator that uses independent temperature control of the MEA to prevent condensation, the system enables stable, uninterrupted recordings for weeks. We show that perturbations in firing rate and functional connectivity following medium exchange are significantly reduced by suppressing evaporation. We demonstrate continuous 35-day recordings of patterned human iPSC-derived neuronal networks with a single medium exchange, revealing the spontaneous emergence and consolidation of spatiotemporal firing patterns during maturation. All design files are provided to facilitate adoption across culturing platforms, enabling uninterrupted longitudinal interfacing with network dynamics for studies of plasticity, chronic pharmacology, and developmental trajectories in individual cultures.

## Introduction

Microelectrode array (MEA) recordings of neuronal cultures enable non-invasive, long-term electrophysiological readout, making them indispensable for drug screening, disease modelling, and fundamental neuroscience (1±4). Longitudinal MEA studies typically rely on periodic recording snapshots interleaved with regular partial medium exchanges to maintain culture conditions (5, 6). However, each exchange transiently perturbs network activity, retains a fraction of altered medium, and poses contamination risks (7, 8). At the low cell densities typical of MEA cultures, the dominant driver of medium deterioration is not metabolic waste accumulation but evaporation-induced changes in ionic concentration (9±11), which is amplified in the case of partial medium exchanges (see Fig. S15). As neuronal cells are highly sensitive to osmolarity shifts (12, 13), particularly in specialised neuronal media with reduced ionic buffering capacity (14), even modest evaporation compromises recording stability and culture health. Eliminating evaporation would allow the interval between medium exchanges to exceed the timescales of longterm plasticity and network maturation— the very processes longitudinal MEA recordings aim to capture.

Existing mitigation strategies face a fundamental trade-off between suppressing water vapour loss and maintaining CO_2_ and O_2_ exchange required for pH buffering and cell viability (9, 15). Diffusion barriers such as sheet membranes (10, 11, 16) or silicone oil overlays (17) reduce but cannot eliminate evaporation, and the latter raises concerns regarding cytotoxicity and media integrity (18, 19). Active humidity control systems can achieve higher suppression, but remain expensive, require dedicated equipment per recording station, and risk condensation damage to sensitive electronics (16, 20). All diffusion-limiting approaches only attenuate evaporation and therefore still require standard half-medium exchange protocols, which lead to elevated, unphysiological ion concentrations as the affected medium remains partially inside the well.

Directly counteracting evaporation by adding purified water requires precise volume feedback control and introduces contamination risk (10, 13). Continuous perfusion systems eliminate ionic drift through constant medium exchange (21±24), but at the cost of complex microfluidic infrastructure, high medium consumption, and shear stress on cells— effectively substituting one source of perturbation for another.

Here, we present a fundamentally different approach: a culture lid with an integrated water compartment positioned above the liquid±air interface, which fully eliminates evaporation at its source by removing the humidity gradient, while preserving gas exchange. Combined with a temperature control system, this design maintains stable culture conditions without requiring medium exchanges, active humidity regulation, or perfusion infrastructure, and is readily scalable to parallel multi-chip recordings with minimal modification to standard workflows.

We show that cultures under the water compartment lid exhibit significantly reduced transient perturbations in firing rate and functional connectivity following medium exchange compared to conventional membrane lids. By suppressing evaporation and its downstream confounds, we achieve what is, to our knowledge, the longest continuous, unperturbed recording of patterned neuronal networks on a high-density MEA— over 35 days with only a single medium exchange. The resulting dataset captures network maturation at unprecedented spatiotemporal resolution, revealing the spontaneous emergence of firing patterns that diversify during development before consolidating into a smaller set of dominant sequences. These dynamics are invisible to conventional periodic recordings. All design files are provided to facilitate adoption across culturing platforms. Our platform enables uninterrupted longitudinal interfacing with network dynamics for previously unfeasible studies of plasticity, chronic pharmacology, and developmental trajectories in individual cultures.

## Results

### Water compartment and temperature control eliminate evaporation while permitting gas exchange

As described by Fick’s law of diffusion, evaporation is driven by the gradient in water vapour concentration (or partial pressure). In a cell culture system with an exposed air-liquid interface, the relative humidity of the air above the medium thus determines the evaporation rate, with dynamics governed by the diffusion barrier. Temperature gradients can further cause condensation on colder surrounding surfaces. Complementary metal±oxide±semiconductor (CMOS) based HD-MEA systems are particularly susceptible to these effects, as active readout electronics on the chip dissipate power and heat up the culture medium. These mechanisms are summarised schematically in Fig. 1a.

**Fig. 1.**
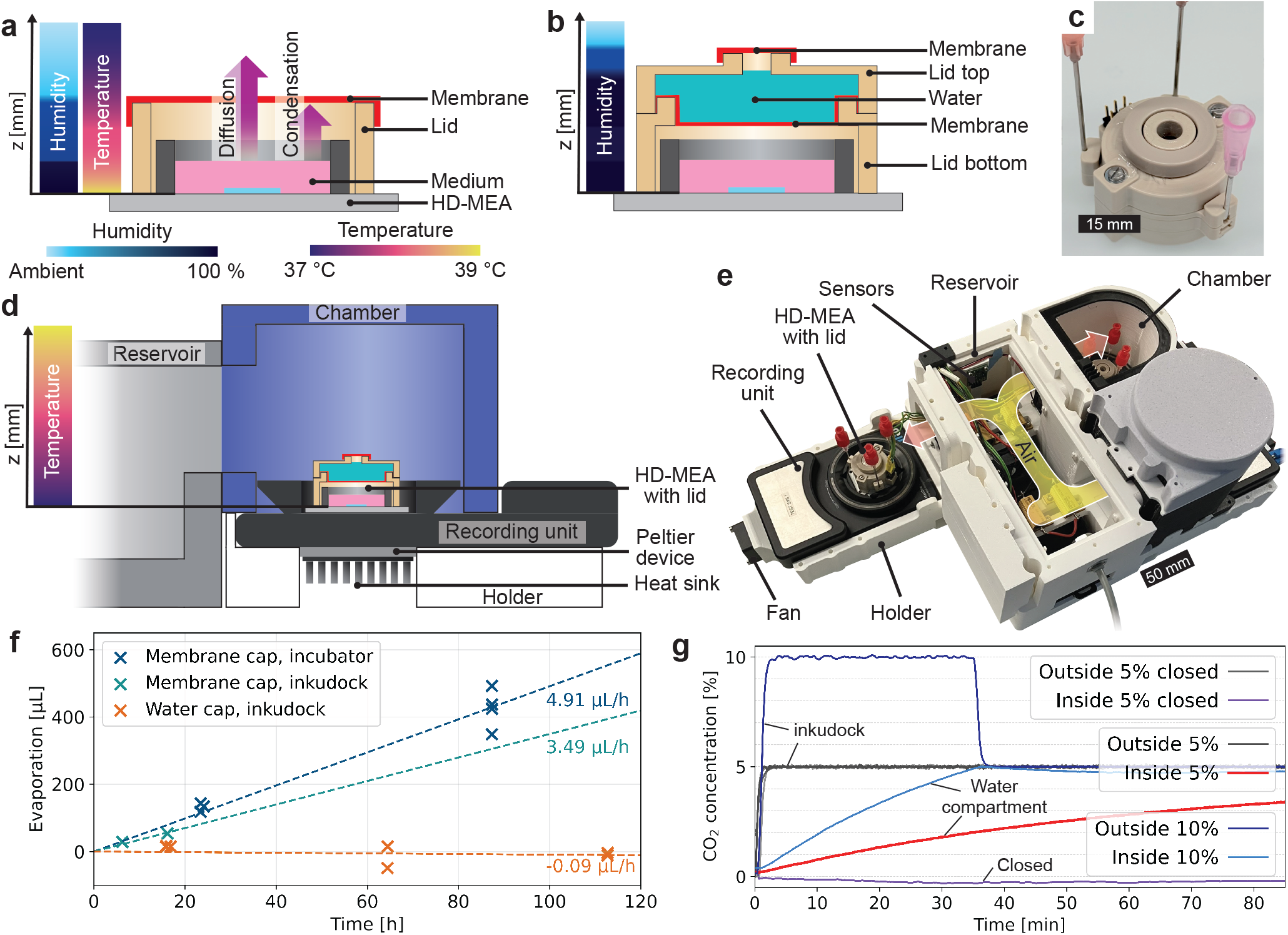
Addressing humidity and temperature gradients eliminates evaporation whilst preserving gas exchange. **a**, Cell culture medium constantly evaporates from dishes due to humidity gradients. In the presence of recording systems, ambient humidity is further limited to prevent damaging electrical components. Additionally, temperature gradients, caused by incubators or power dissipation of active HD-MEAs, lead to an elevated chip surface temperature with respect to its culturing environment. This raises evaporation levels and leads to condensation when common plastic or gas-permeable membrane lids are employed, which enclose the chip to reduce evaporation and contamination risk. **b**, Cross-section of a culture lid with 2 gas-permeable membranes (red) that enclose a water compartment (cyan), eliminating humidity gradients. **c**, Depiction of the manufactured lid with a top part fixed with screws. The lower air compartment and the water compartment are hermetically sealed with three O-rings. **d**, A custom incubator setup inkudock, that controls the chip temperature to prevent condensation. Cross-section of the design. The central reservoir resembles a standard incubator. A chamber is attached and encloses the chip with the lid. The recording unit is exposed to the ambient environment, therefore enabling easy temperature control by heating and cooling with a thermoelectric Peltier device. **e**, A photo showing inkudock. Up to four chambers can be attached to one reservoir, distributing the airflow to all attached chips. **f**, Evaporation from HD-MEAs during recording is quantified in different setups by weighing the chips before and after. A linear fit is applied to determine the evaporation rate. When the temperature gradient is reversed by placing the chip with a single membrane lid (membrane cap) in inkudock, evaporation is reduced by about a third. Applying the water compartment lid (water cap) effectively eliminates the evaporation. **g**, A sensor dummy that matches the dimensions of an HD-MEA but carries a sensor board for humidity, ambient temperature, and CO_2_ concentration (see Supplementary Information A.3.1) is used to measure the gas exchange. A lid where the lower membrane of the water compartment is replaced with solid material shows no increase in CO_2_ after a step to 5 % in the inkudock, proving airtightness of the design. In contrast, the water cap shows CO_2_ transmission with a *τ* of about 75 min, averaged over 3 measurements. The time required for saturation at 5 % can be shortened by stepping the ambient CO_2_ concentration to 10 % for 35 min.

The proposed system introduces a water compartment positioned above a vapour-permeable membrane, creating an interface with 100 % humidity on the upper side. This compartment thereby serves as the buffer interface to the low-humidity surroundings, isolating the culture medium below. To prevent depletion of the water compartment itself, it is sealed with a second membrane. The water layer creates a hermetically closed system. The working principle is illustrated in cross-section in Fig. 1b, with a manufactured prototype shown in Fig. 1c.

To eliminate condensation, the temperature gradient must be balanced or reversed such that the medium and chip surface are cooler than the lid surfaces. We present a custom incubator (‘inkudock’, Fig. 1d-e), where a central reservoir serves as a standard incubator and individual chambers are attached that enclose an HD-MEA mounted in the commercial recording unit (MaxOne+ and MaxOne Recording Unit, Maxwell Biosystems AG, Zurich, Switzerland). This seals the inside of the incubator from ambient conditions whilst keeping the bulk of the recording unit exposed to the outside, which enables independent temperature control. As the chip is strongly thermally coupled to the recording unit, the lid can be heated via the reservoir temperature without strongly affecting the chip temperature. Alternatively, the water compartment can be actively heated to invert the temperature gradient when placed in a standard commercial incubator (see Supplementary Information A.1.3).

To validate the system’s performance, we first quantified evaporation by weighing HD-MEAs before and after continuous recording under different configurations (Fig. 1f). To achieve physiological conditions for the cells in inkudock, the HD-MEA surface temperature was set to 38.5 °C, resulting in a medium temperature of 35.5 °C at a chamber temperature of about 43 °C (for calibration procedure see Supplementary Information B.3). The temperature in a commercial incubator with a passive water bath (CB170, BINDER GmbH, Tuttlingen, Germany) was set to 35 °C. Placing a chip with a common single-membrane lid (see Methods) in inkudock reduced evaporation by approximately 30 % compared to the incubator. Evaporation rate without recording in the incubator ranges between 0.8 (see Fig. S14) and 2.5 µL/h (16). The water compartment lid effectively eliminated evaporation, with one negative weight change attributed to condensation on the culture medium due to suboptimal temperature sensor placement.

Gas exchange through the water compartment was tested using a validation chip with dimensions matching the employed HD-MEAs, but equipped with a sensor board for humidity, temperature, and CO_2_ concentration (see Fig. S12). Fig. 1g shows the CO_2_ concentration step response. A control measurement with the lower membrane replaced by solid material confirmed hermetic sealing of the remaining design. With the water compartment, CO_2_ transmission exhibited a time constant *τ* of approximately 75 min with no liquid in the well. To accelerate saturation, we applied an initial step to 10 % CO_2_ for 35 min, before returning to 5 %. Additional tests with CO_2_-saturated water indicated that transmission from the water compartment to the lower air gap dominates the saturation dynamics. At the cell densities typical of MEA cultures, these CO_2_ equilibration dynamics are expected to maintain pH buffering.

### Transient changes in network activity caused by medium exchange are suppressed through evaporation-suppressing lid

Patterning neuronal networks in microfluidic devices prevents cell migration and thus forms an ideal platform for longitudinal recordings (26, 27). We used patterned networks to assess the effects of reduced evaporation on electrophysiology and network dynamics. PDMS membranes confine cell bodies in seeding wells, which are connected through microchannels. We applied them to HD-MEAs and selected the microchannel regions for continuous recording of action potentials (overview in Fig. 2a, for details see Methods). The impact of medium exchange on the firing rate has been documented in literature (6, 28, 29) and is accounted for in MEA recording protocols (30). To investigate whether evaporation-induced osmolarity shifts contribute to these changes, we compared cultures equipped with a membrane-sealed lid versus our water compartment lid, assuming previously determined evaporation rates. We continuously recorded the cultures in inkudock and performed a medium exchange every 3 to 4 days. Networks cultured with a conventional membrane lid exhibited significantly greater transient changes in the relative firing rate immediately following medium exchange when performing a linear fit (Fig. 2b-d).

**Fig. 2.**
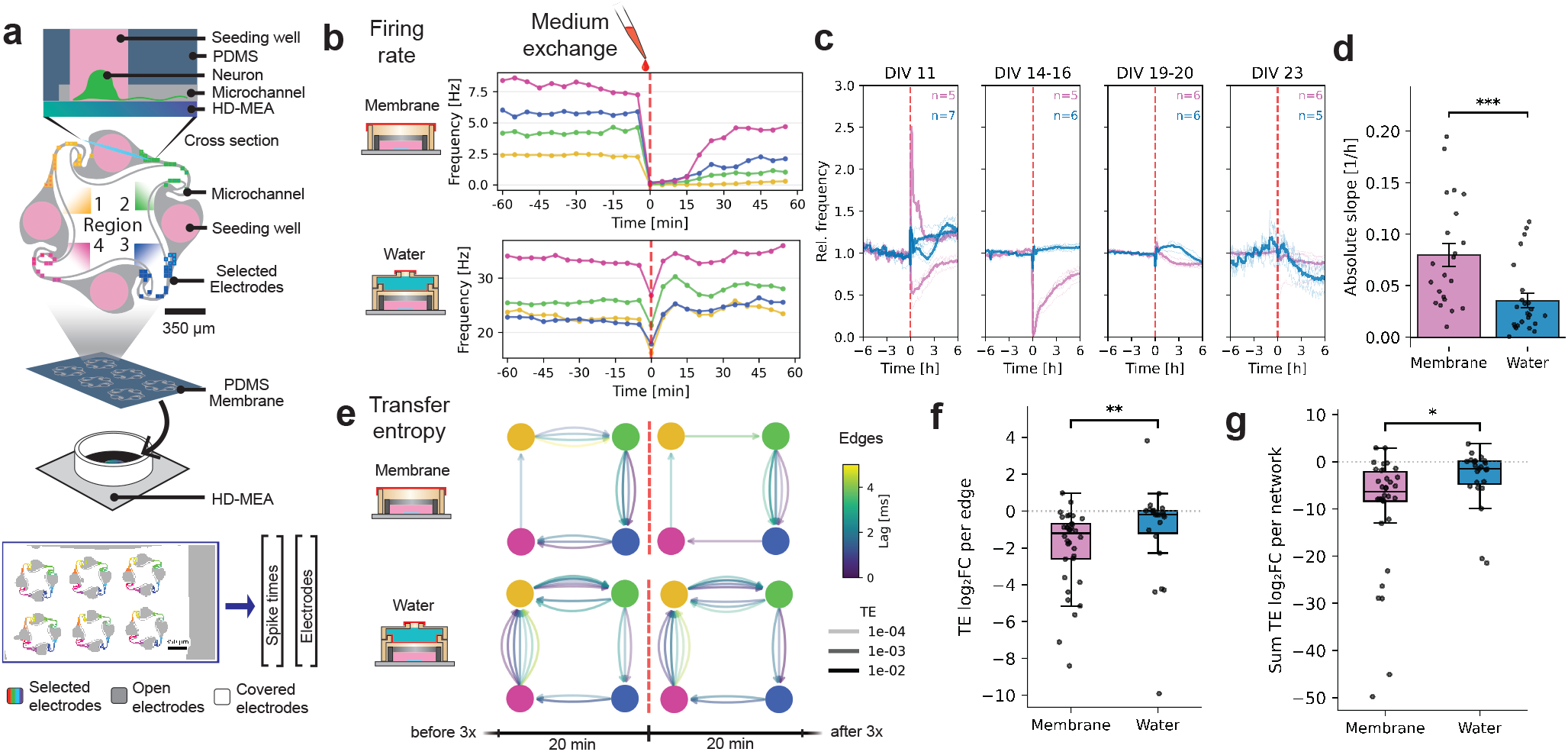
Comparison of the impact of medium exchange on network activity. **a**, Compartmentalised networks were formed on top of an HD-MEA with a two-layer PDMS membrane, which restricts cell bodies from migrating from the seeding well into the microchannels (25). Electrodes were selected inside the microchannels with an impedance scan and summarised into 4 regions. Spikes from up to 6 networks in parallel were simultaneously recorded using the MaxLab Live application programming interface (API). **b**, A chip with a single-membrane lid and one with the water lid were placed inside inkudock and recorded continuously. Every 3 to 4 days, half of the culture medium was exchanged with fresh medium. The firing rates of the 4 regions for one example network per condition 1 h before and after the exchange are shown, averaged across 5 min. **c**, The firing rate of several networks on different MEAs, divided by the mean firing rate in the 6 h before an exchange are shown for different DIV. The average across the networks on the same MEA is shown as thick lines. **d**, The slope of a linear fit on the relative rate change in the first 4 h after exchange was compared across the conditions. Mann±Whitney *U* testing shows significantly less impact of medium exchange on the activity with the water compartment lid (20 unique networks; *p <* 0.001; linear mixed model with network identity as random intercept confirmed negligible between-network variance across DIVs). **e**, Transfer entropy between network regions was calculated as described in Methods. Example networks 20 min before and after the exchange are shown for both conditions. Only networks with a weakly connected component of at least 3 nodes were used for further analysis. The total analysis comprised 25 unique networks with 81 network-exchange observations (54 membrane, 27 water). **f**, Persistence of edges across medium exchange was assessed. Stable edges were defined as connections with TE ≥ 1 × 10^−4^ at the same lag across all three 20 min windows before exchange. The mean log_2_ fold change of transfer entropy per stable edge is shown (disappearing edges were floored to TE = 1 × 10^−6^). Networks with the water compartment lid showed significantly lower changes (Mann±Whitney *U* ; *p <* 0.01). The number of stable edges was 175 and 113 for the membrane and water compartment lid, respectively. **g**, The fold change summed across a network also showed higher persistence with the water compartment lid (linear mixed model with network as random intercept; *p <* 0.05).

Functional connectivity within a network is a common read-out for assessing circuit-level dynamics (31, 32). We therefore tested whether medium exchange altered directed information flow beyond global changes in firing rate. Effective connectivity was quantified using transfer entropy within a 5 ms time window, computed from 20 min segments of spike data. Fig. 2e shows the transfer entropy as directed edges with different lags. Stable directed edges were defined as connections with TE ≥ 1 × 10^−4^ at a consistent lag across all three pre-exchange windows. Edges were tracked across the exchange by computing the log_2_ fold change of their mean TE with the TE at the same lag after exchange (Fig. 2f,g). The mean fold change per edge showed a roughly 2.5-fold stronger reduction with the membrane lid (Mann±Whitney *U, p <* 0.01). Also, when summed across a network, the membrane lid showed stronger reorganisation (linear mixed model with network as random intercept, *p <* 0.05). Medium exchange thus induced a more pronounced reconfiguration of directed functional interactions for the membrane lid in the presence of substantial evaporation. This indicates a transient displacement of the network from its ongoing dynamical regime, affecting information-theoretic readouts.

Notably, since evaporation with the membrane lid in inkudock is substantially lower than the rates expected during continuous recording in a commercial incubator, the observed effects are a conservative estimate of the medium exchange impact under typical recording conditions.

### Persistent network activity can be continuously recorded for weeks without intervention

With our system, recordings that were previously limited to a few hours due to evaporation-induced changes in firing rates can now be performed for days, with longer intervals for medium exchange (also see comparison of culture moved between conditions in Fig. S23). This enables the characterisation of network maturation without the confounding effects of medium exchange. We performed continuous recordings of two HD-MEAs equipped with water compartment lids for 35 days (see Fig. 3 for electrophysiology, environmental parameters are shown in Fig. S17). Recording was started at 7 DIV following a full medium exchange, and interrupted after 3 weeks by a single exchange, where half of the culture medium was replaced with fresh medium. Both chips exhibited remarkably similar developmental trajectories, with network activity emerging after a few days, as is expected from these cultures (33, 34). Activity levels rose progressively before plateauing between 14 and 21 DIV, after which firing rates declined on both chips. Notably, the proportion of active units decreased less dramatically than the overall firing rate, indicating that many electrodes continued to record sparse activity even as network-wide firing diminished. This decline could be attributed to nutrient depletion, as the exchange of culture medium initiated a slow recovery. Glucose measurements showed a decrease by 20 % in 14 days, while protein electrophoresis and UV-vis spectroscopy indicated stable levels of protein content (see Supplementary Information B.5). Another potential cause could be an empty water compartment leading to residual evaporation, supported by the similar trajectories of both MEAs. If nutrient depletion or waste accumulation were responsible, one would expect different onset times, as the two cultures differ slightly in cell count and medium condition and therefore in metabolic demand.

**Fig. 3.**
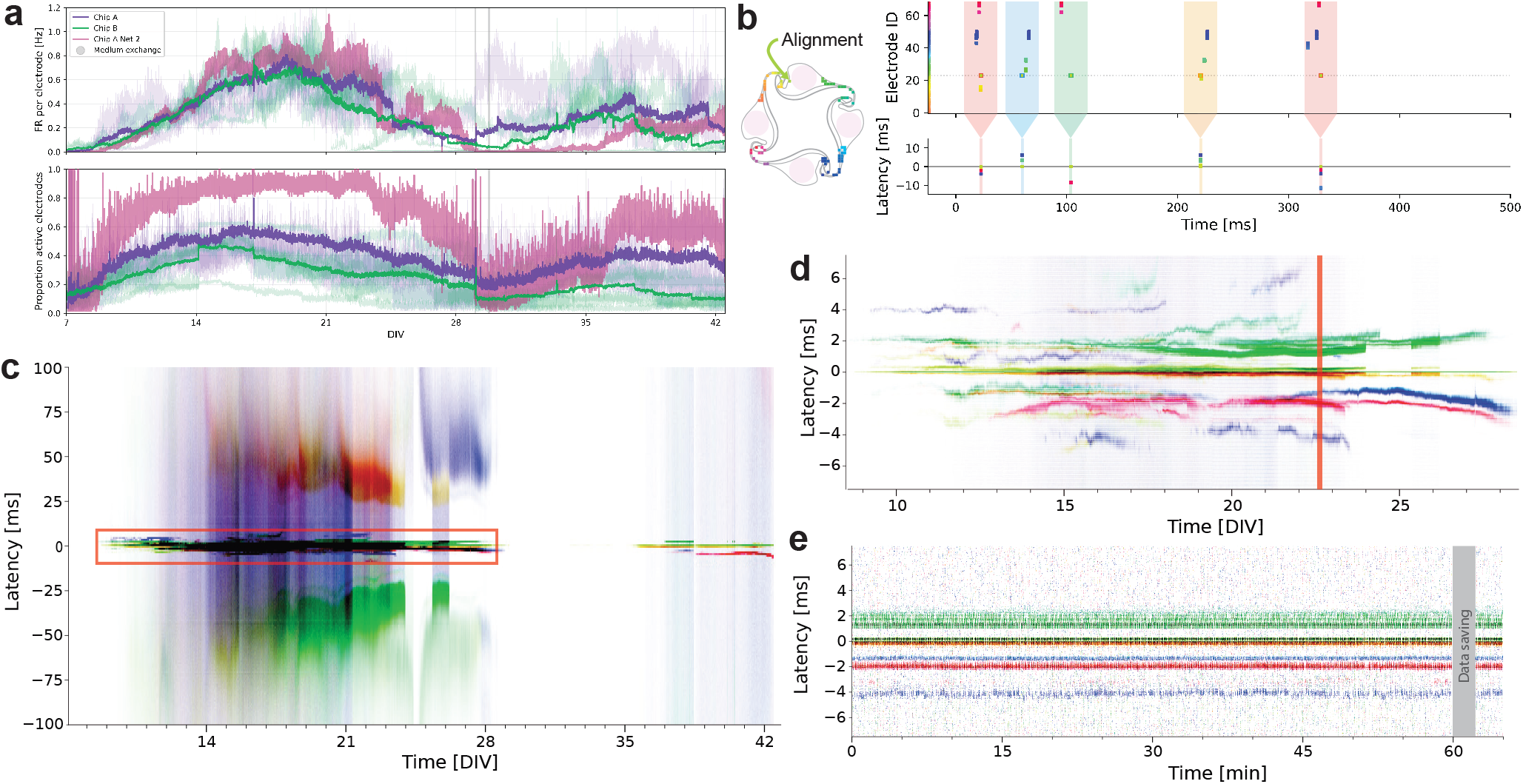
Network activity of two HD-MEAs with a water compartment lid is continuously recorded for 35 days. **a**, Firing rate per electrode and proportion of active electrodes over time. Networks are considered active if their average firing rate lies above 0.1 Hz. The HD-MEAs contain 6 independent networks each (faint lines, with network 2 from chip A highlighted in pink). The recording was started at 7 DIV, with a full medium exchange prior to the recording and an exchange of half the medium after 3 weeks. The water compartments were refilled after 10 days. Spikes were recorded for 1 h followed by a break of 2 min for data saving. The firing rate decreased after 21 DIV on both chips, but recovered after the medium exchange. **b**, Spike-time-triggered raster plots (STTRPs) are generated to visualise network activity over days. The most active electrode in a network is used for alignment, and spikes in its vicinity (coloured windows in the upper axis, here for ±15 ms) are plotted with their respective latencies at the spike-time of the alignment spike (lower axis). **c**, Full STTRP of network 2 from chip A with a maximum latency of ±100 ms, binned to 30 min. Spikes are colour-coded following panel b, with a firing frequency of 2 Hz leading to saturation of a pixel with the electrode colour. After 14 DIV, synaptic activity formed bands of consistent latencies at ±50 ms. Over a time course spanning about 2 weeks latency decreased until the firing at this relatively stable latency abruptly disappeared. **d**, Zoom-in on 8.5 to 28.5 DIV and a latency of ±10 ms (red rectangle in panel c). Sequential spikes across the whole network are visible, presumably relying on fast synaptic transmission between the network nodes. Several bands of activity with consistent latency and low probability occur, especially between 14 and 24 DIV. **e**, Zoom-in to 1 h on 22.5 DIV (red bar in panel d). The activity is remarkably stable during the conventionally applied recording duration of tens of minutes.

We performed a detailed analysis of the spatiotemporal network dynamics over time on network 2 from chip A (firing rate shown in Fig. 3a), where we acquired about 800 million spikes across 35 days. For visualisation, we aligned the spontaneous network activity to its most active electrode and investigated the relative latencies of spikes on the remaining network electrodes over time with a spike-time-triggered raster plot (STTRP, see Fig. 3b). When spikes reliably occur at a similar latency over time, a horizontal band is visible in the STTRP, suggesting a strong connection between the neurons recorded on those electrodes. At long latencies, synaptic activity bands appear around ±50 ms, indicative of polysynaptic transmission from the alignment to the observation electrode (35±37). Zooming in to shorter timescales (±10 ms) reveals circular propagation patterns consistent with the directional recurrent network architecture. As activity begins to decline, firing latencies increase, suggesting slowed conduction or weakened synaptic transmission. An additional network is presented in the Supplementary Information B.8 with Fig. S24 and Fig. S25.

### Diversity of spontaneous firing patterns increases during network maturation

Persistent sequences of spontaneous activity are a frequently studied target, due to their assumed importance for memory formation and retention (38, 39). Analysing their trajectories through continuous recording represents one of the many potential benefits of the system presented here. To quantify the emergence and reinforcement of spontaneous firing patterns during network maturation, we developed a clustering pipeline for low-latency sequences of activity (Fig. 4a-c). To reduce the electrode dimensionality, spikes recorded within a window of 1.5 ms on multiple electrodes belonging to the same region of the microstructure corresponding to the underlying microchannel geometry were summarised into a region event (for details see Methods). Region-specific firing sequences were then aligned to the first region event (on the region with the lowest ID), windowed to ±10 ms around this reference, and converted into matrices. A density-based clustering (HDB-SCAN, see Methods) was performed on the first 5 principal components and visualised via UMAP, showing clear cluster separation for most patterns (Fig. 4d). Example sequences from the 4 most prominent clusters, which stand for a consistent activity pattern, demonstrate low intra-cluster variability. All patterns sorted by the first principal component are provided in Fig. S26. Fig. 4e shows when and how often specific patterns occurred during the experiment. Different phases can be identified in Fig. 4f-g, during which the diversity of patterns changed. The number of concurrently active patterns increased steadily until it plateaued at 17 DIV. The first stable firing patterns emerged around 10 DIV, with multiple low-frequency sequences coexisting as early as 12 DIV up to 24 DIV alongside an increasingly dominant cluster. This could indicate cell maturation and early synapse formation, which is reported after 2 to 4 weeks *in vitro* (40, 41). After 24 DIV, the network transitioned to a state with fewer simultaneously active patterns, indicating more stable synaptic connections (42, 43). Following the period of near-silence, activity re-emerged at 36 DIV, with single prominent pattern. As some patterns are shared between both periods, a reactivation of persisting pathways is more likely than novel connections.

**Fig. 4.**
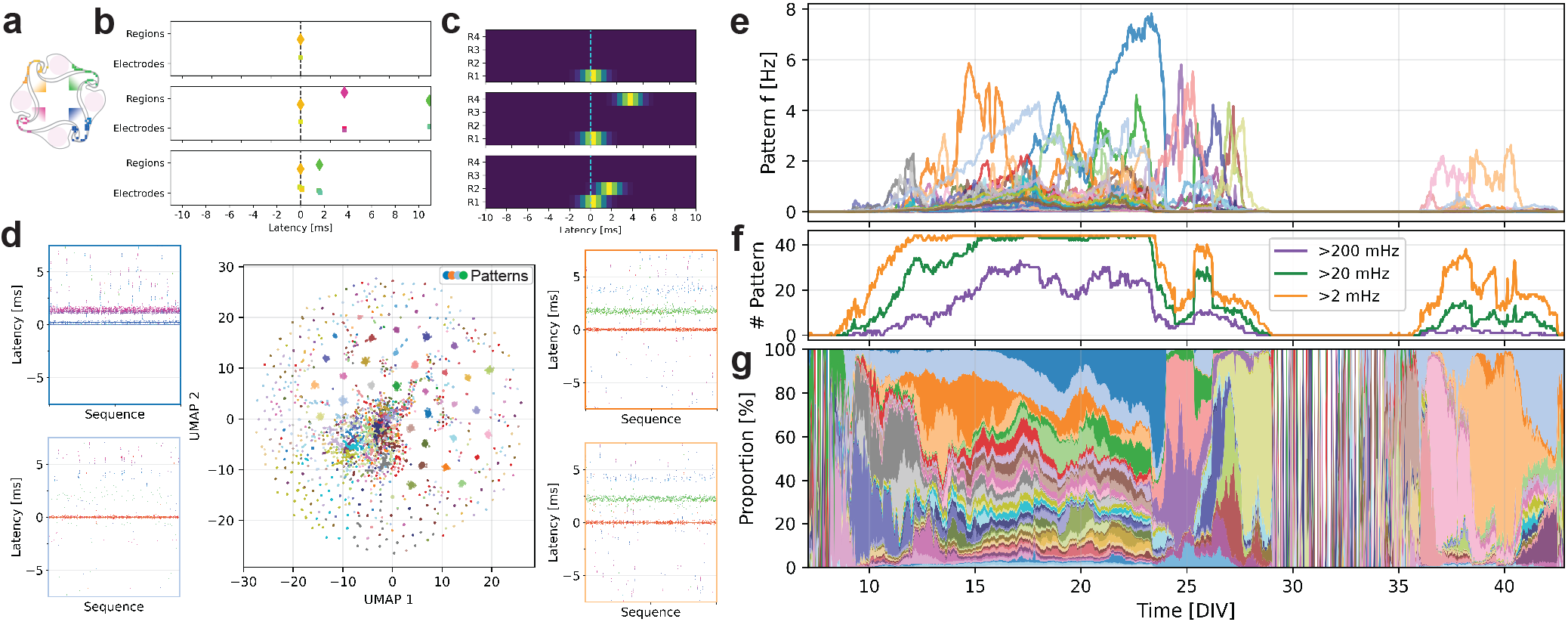
Development and reinforcement of spontaneous firing patterns during maturation. **a**, Network electrodes were aggregated into 4 regions corresponding to the microchannels. A colour code was applied, mapping the hue circle onto the angular rank of the electrodes with gaps for the wells. **b**, The most active electrode was selected for each region and firing sequences, with at least one spike detected within ±10 ms on another electrode, were extracted. The dimensionality of the firing sequences was reduced for clustering. When more than 2 spikes occur in the same region within 1.5 ms, they were registered as a region event (diamonds). A few examples are shown next to the original spike data by electrode, following the colour code in **a**. About 25 million sequences were identified. **c**, Sequences were aligned to the first region event (in the region with the lowest ID) and transformed into spatiotemporal matrices spanning ±10 ms with a Gaussian kernel as described in Methods. **d**, PCA was performed on the flattened matrices. Clustering was performed using HDBSCAN on the first five principal components, yielding 44 recurring patterns. The 2D UMAP shows the cluster distribution with 500 example sequences from the 4 most frequent patterns (for all sequences see Fig. S26). **e**, The frequency of the patterns was tracked over the whole experiment duration and aggregated into bins of 1 h. The colour code is selected according to the UMAP. Stable patterns emerged around 10 DIV. The most frequent pattern peaks at more than 10 Hz. **f**, The number of patterns surpassing 3 different frequency thresholds is shown over time. When looking at the pattern diversity, the number of patterns occurring with a frequency above 20 mHz plateaus at about 15 DIV and drops rapidly from 23 to 24 DIV. **g**, The distribution of patterns is shown by proportion. After 36 DIV, single patterns dominate the distribution.

Prior studies have described a rich repertoire of evolving burst patterns during maturation of dissociated cortical cultures, with pattern diversity highest while synaptic connections are still forming (44). Periodic recordings have shown that synchronised activity emerges progressively and stabilises once mature (45), and continuous perfusion-based recordings confirmed that activity remains broadly stable in the absence of external perturbation (22). These observations were largely limited to population-level burst statistics, as passive low-density MEAs sample only a fraction of active units in large, randomly connected networks. The patterned networks and HD-MEA recordings employed here extend this view to single-spike spatiotemporal sequences, enabling the tracking of individual firing patterns across weeks of uninterrupted maturation.

## Discussion and Conclusions

With the advancements in cellular models and MEA technology, an easily usable, ready-to-scale solution eliminating evaporation-induced changes in ionic concentration is required for higher-quality assays and longitudinal studies.

We developed a hermetically sealed cell culture lid with a water compartment interface that eliminates evaporation of cell culture medium whilst maintaining gas permeability. Combined with integrated temperature sensing and a custom incubator that prevents condensation through independent temperature control, the system enables stable weeks-long MEA recordings without medium exchange. We were able to track the maturation process and firing pattern diversity.

To improve usability, membrane protection against rupturing, overpressure and refill access ports, and an integrated, wireless temperature control of the water compartment for use in commercial incubators could be added. Furthermore, active humidification of inkudock, a smart refill system, or a humidified chamber above the second membrane, can prevent an empty water compartment. The presented lid design is easily adaptable to other culturing formats, such as multiwell plates.

Our platform addresses a fundamental limitation in longterm electrophysiology by providing the environmental stability necessary for reliable effect attribution in studies of network maturation, plasticity, and chronic pharmacological interventions. By enabling continuous recordings without medium exchange, the system reduces media consumption and manual intervention whilst improving culture sterility through hermetic sealing and relaxing constraints on incubator humidity. Notably, we demonstrated compatibility with human iPSC-derived neurons, which represent a highly relevant model system for studying neurodevelopmental processes and disease-associated phenotypes yet are particularly sensitive to environmental perturbations (46).

Continuous monitoring offers several advantages over periodic recording snapshots. It captures transient phenomena that would otherwise be missed, and reduces handling-induced variability. It also facilitates interfacing cultures with computational models for closed-loop experiments. Finally, fewer biological replicates are needed to achieve the same statistical power, as within-culture variance is reduced. Together, these improvements enable more robust and efficient investigation of long-term network dynamics.

## Methods

### Hardware description

#### Water compartment lid

The lid comprises six CNC-milled components made of polyether-ether-ketone (PEEK) assembled with epoxy adhesive and screws. Two membranes positioned above the chip well create a water-filled compartment with an air gap. Three bent stainless-steel needles enable liquid handling during operation. A laser-cut copper plate electrically grounds the needles via screw contacts, preventing noise induction. A PT1000 temperature sensor integrated into the bottom component monitors chip surface temperature. Airtight sealing is achieved through epoxy bonding along the glue ridge and compression of o-rings at the membrane and chip interfaces. Detailed fabrication instructions, including a reduced version without fluidic inlets, are provided in Supplementary Information A.1; design files are available in the repository listed in the data availability statement.

#### Custom incubator

All experiments were performed in a custom incubator (inkudock) consisting of a central reservoir and up to four attachable chambers, all 3D-printed from polylactic acid (PLA). The reservoir contains a sensor board (47), resistive heaters, a CO_2_ inlet, and circulation fans. Each chamber is mounted atop a MaxOne recording unit, which sits on a 3D-printed holder equipped with a Peltier device and heat sink for temperature control. Electronics were adapted from the inkube system (13), with temperature and CO_2_ control for the reservoir and four independent temperature control loops for the recording units. Sensor signals are routed through a custom shield on the Arty Z7 development board (Digilent, Pullman, WA, USA) to a system-on-chip (XC7Z020-1CLG400C, Xilinx, San Jose, CA, USA), which then transmits control signals through the shield to a power board for amplification. Modifications include an updated shield with PT1000 readout circuitry and a revised power board with more powerful single-ended amplifiers for reservoir temperature and humidity control. Hardware design files, derived from the inkube system (13), are released under the Creative Commons Attribution-ShareAlike 4.0 International licence (CC BY-SA 4.0; https://creativecommons.org/licenses/by-sa/4.0/), see data availability statement. Assembly instructions are provided in the Supplementary Information A.2.

#### Sensor dummy chip

A sensor dummy chip with dimensions matching the HD-MEA was CNC-manufactured from PMMA. The dummy incorporated an inkusense sensor board (47) equipped with humidity, temperature, and CO_2_ sensors, positioned on top of a dummy well that can be filled with liquid. Cables were routed through the bottom of the assembly and hermetically sealed with epoxy (see Supplementary Information A.3.1).

#### Sensor ring

A custom sensor ring mounted beneath the conventional single-membrane lid contains a PT1000 temperature sensor pressed to the chip surface. The ring was fabricated using LCD resin 3D printing. Details are provided in the Supplementary Information A.3.2.

### Cell culturing

#### Chip preparation

High-density microelectrode arrays were prepared with polydimethylsiloxane (PDMS) microstructures as described previously (48). Briefly, the chip surface was coated with poly-D-lysine (PDL), and a PDMS membrane was cut from a wafer and positioned onto the dried surface using tweezers. The MEA well was filled with phosphate-buffered saline (PBS) and desiccated to remove air bubbles, then PBS was replaced with culture medium before cell seeding. The two-layered PDMS microstructures, manufactured by Wunderlichips GmbH (Zurich, Switzerland) using soft lithography, consisted of open-top nodes for cell bodies connected by enclosed microchannels. The microstructures employed 4-node recurrent designs that promote directional axonal growth (25), with 6 full networks fitting onto the chip sensing area.

#### Culture preparation

Early-stage Ngn2-induced neurons derived from human iPSCs (33) were obtained from Novartis (Basel, Switzerland), thawed and differentiated using neurobasal differentiation medium (NBD+): Neurobasal Plus (A3582901) with GlutaMAX (35050-061) and Pen-Strep (15070-063), supplemented with B-27 Plus (A3582801), N2 (17502-048), BDNF (10 µg mL^-1^, 450-10) and GDNF (10 µg mL^-1^, 450-02, all Thermo Fisher Scientific, Waltham, MA, USA), as previously described (34). For suspension seeding (long-term recordings), cells were plated at 150,000 cells per chip in NBD supplemented with 5 µg/mL laminin (L2020, Sigma Aldrich, St. Louis, MO, USA) and 2 µg/mL doxycycline (631311, Takara Bio USA, Inc., Mountain View, CA, USA). Complete medium exchanges were performed at 60 min and 7 days post-seeding. For spheroid seeding (comparison experiments), cells were aggregated at 100 cells per spheroid using AggreWell^*T M*^ 400 plates (34415, StemCell Technologies, Vancouver, Canada) following the manufacturer’s protocol. Spheroids were seeded after 1 day into medium supplemented with doxycycline and laminin, with full exchange at 7 days post-plating. Doxycycline and laminin supplementation was maintained for the first 7 days after plating in both protocols.

#### Membrane lid

MEAs were kept under a standard single-membrane lid outside of experiments and during controls. We used an open-source lid, kindly provided by the University of California Santa Cruz, based on the original by Potter *et al*. (10), optimised for the MaxOne chip geometry with a tighter fit (see Supplementary Fig. 3 in (21); design available on GitHub as membrane_ring.stl (49)). The outer holder ring was sealed with a fluorinated ethylene propylene membrane sheet of 12.5 µm thickness (ALA MEA-MEM-SHEET, ALA Scientific Instruments Inc.).

### Electrophysiology

#### Electrode selection

Due to the 1024-channel recording limit of the MaxOne HD-MEA, electrodes were preselected based on PDMS microstructure placement. An impedance scan was performed as described in previous work (50). A threshold was applied to the impedance map to identify PDMS-covered regions and distinguish them from open electrode sites. Electrodes within the open regions in the microchannels connecting the wells were selected for analysis, with some selections expanded to include all electrodes from one entire network.

#### Spike data

Spike detection was performed using the MaxLab Live Python API (Maxwell Biosystems AG, Zurich, Switzerland) with detection thresholds of 4.5 standard deviations for exchange experiments and 5 standard deviations for longterm recordings. Neural activity was recorded in continuous chunks of 1 (long-term) or 2 hours (exchange). Time delays between chunks interrupted by medium exchange were calculated from file generation timestamps, as the spike timing resets upon hardware restart.

### Network activity analysis

#### Spike data dimensionality reduction

For transfer entropy and pattern clustering, the electrode dimensionality was reduced. Electrodes located beneath the same microchannel record largely redundant activity; dimensionality reduction was therefore applied to obtain a single representative event. Each electrode was assigned to one of four regions based on its angular position relative to the network centre, yielding four quadrants corresponding to the microchannels. Each of the regions comprises about 30 to 50 electrodes. Within each region, spikes occurring within a 1.5 ms window were merged into a single event with the mean spike time, while single spikes were omitted. As the typical travelling time of an action potential through the microchannel is about 1 ms this time window prevents duplicate detection, while still allowing separation of 2 consecutive spikes.

#### Transfer entropy

Pairwise transfer entropy (TE) was computed between all region pairs of the membrane and water lid conditions at lags up to 5 ms using 0.5 ms bins. Spike data was aggregated over 6 non-overlapping 20 min windows spanning a 2 h period centred on each medium exchange event. Each TE value at a certain lag was considered as a directed edge from source to target in the network graph, with the graph nodes corresponding to the 4 regions. Statistical significance of each inferred link was assessed using surrogate testing with 100 surrogate time series. For each network, resulting *p*-values were corrected for multiple comparisons using the Benjamini±Hochberg false discovery rate procedure, and links with adjusted *p <* 0.05 were considered significant.

To quantify network reconfiguration induced by medium exchange, stable functional edges were identified as directed connections with raw TE ≥ 1 × 10^−4^ present at the same lag in all three pre-exchange windows. Contiguous lags were grouped up to 3 bins (1.5 ms), with the requirement that at least one lag within the band exceeded the threshold. For each stable edge, the mean TE across the three post-exchange windows was compared to the mean TE across the three pre-exchange windows by computing the log_2_ fold change, using a floor value of 1 × 10^−6^ to handle cases where post-exchange TE dropped to zero. Per-network summaries in-cluded the mean fold change per edge and the cumulative (summed) fold change across all stable edges.

#### Spike time triggered raster plots

To visualise network activity over time, relative latency of spikes was obtained by aligning spontaneous activity to the spike timing of a single trigger electrode (13). STTRPs were generated using Python. The trigger electrode was selected based on the highest spike count, retaining only the last trigger spike when multiple occurred within one analysis window. Spikes were aligned by subtracting both the nearest preceding and following trigger times, then windowed to ±10 ms relative to triggers. Circular colour coding was applied based on electrode angular position from array centre. Visualisation used 0.1 ms latency bins, 5 s time bins, and colour saturation at 10 spikes per bin.

#### Pattern detection and clustering

Sequences were identified by detecting at least two region events within a time window of 20 ms in the relative latencies aligned to the most active electrode in a region. Each sequence was aligned to the first event in the region of the lowest index and transformed into a matrix of 40 time bins of 0.5 ms spanning ±10 ms around this reference. Region events were convolved with a Gaussian kernel of *σ* = 0.75 ms to account for jitter of transduction latencies and avoid dissimilarity between sequences with shared structure. Principal component analysis (PCA) was applied to the flattened matrices; the first five components were retained, preserving 64 % of the variance.

Clustering was performed on the five-dimensional PC space using HDBSCAN (51), a density-based hierarchical clustering algorithm that infers cluster count from the data density structure, with a minimum cluster size of 2000 sequences and a minimum of 50 sequences required to define a core point of a cluster. Cluster assignment for the full dataset was performed by nearest-centroid projection from a representative subsample of 500,000 sequences used to fit the model. A total of 44 clusters were identified. For visualisation, a 2D UMAP embedding (52) was computed on an equal-per-cluster sub-sample of up to 5,000 sequences per cluster, ensuring that dominant clusters did not disproportionately determine the embedding layout.

## ACKNOWLEDGEMENTS

This research was supported by ETH Zurich, and the Swiss National Science Foundation (SNSF) [project numbers 182779 and 10001282]. We would like to thank Kateryna Voitiuk from the University of California Santa Cruz, for supplying us with membrane lids for the MaxOne HD-MEAs. We would like to thank Christian Frei from the Laboratory of Biosensors and Bioelectronics for his help with protein electrophoresis.

## Data and material availability

Manufacturing files and assembly instructions for the lid and inkudock are available at https://doi.org/10.3929/ethz-c-000797812 under the Creative Commons Attribution±ShareAlike 4.0 International licence (CC BY-SA 4.0). Electrophysiology data and analysis scripts are available at https://doi.org/10.3929/ethz-c-000798160.

The software for inkudock is available on GitHub

https://github.com/maurer-lbb/inkube_software.git.

## CRediT author contributions Benedikt Maurer

Conceptualisation, Methodology, Software, Validation, Formal analysis, Investigation, Data Curation, Writing - Original Draft, Writing - Review & Editing, Visualisation, Supervision. Fabio Fischer: Methodology, Validation, Investigation, Writing - Review & Editing, Visualisation. Giulia Amos: Software, Validation, Investigation, Writing - Review & Editing, Supervision. Vaiva Vasiliauskaitė: Methodology, Software, Validation, Formal analysis, Writing - Review & Editing. János Vörös: Conceptualisation, Methodology, Writing - Review & Editing, Supervision, Project administration, Funding acquisition.

