## Supplementary Information for "A water compartment cell culture lid enables stable longitudinal recording of neuronal networks in vitro"

### Supplementary Note A: Hardware description

#### A.1. Water compartment lid.

***A.1.1. Water compartment lid with fluidics and sensor.*** The water compartment cap, illustrated in Fig. S1, comprises six custom-designed components (C1-C6 in Fig. S1), each CNC-milled from either polyether-ether-ketone (PEEK) or polymethyl methacrylate (PMMA), depending on sterilisation and functional requirements. The components were assembled using Araldite 2011 (Huntsman Corporation, The Woodlands, Texas, USA), a two-component epoxy adhesive selected for its mechanical strength and chemical stability. A comprehensive list of all components is available in Table 1.

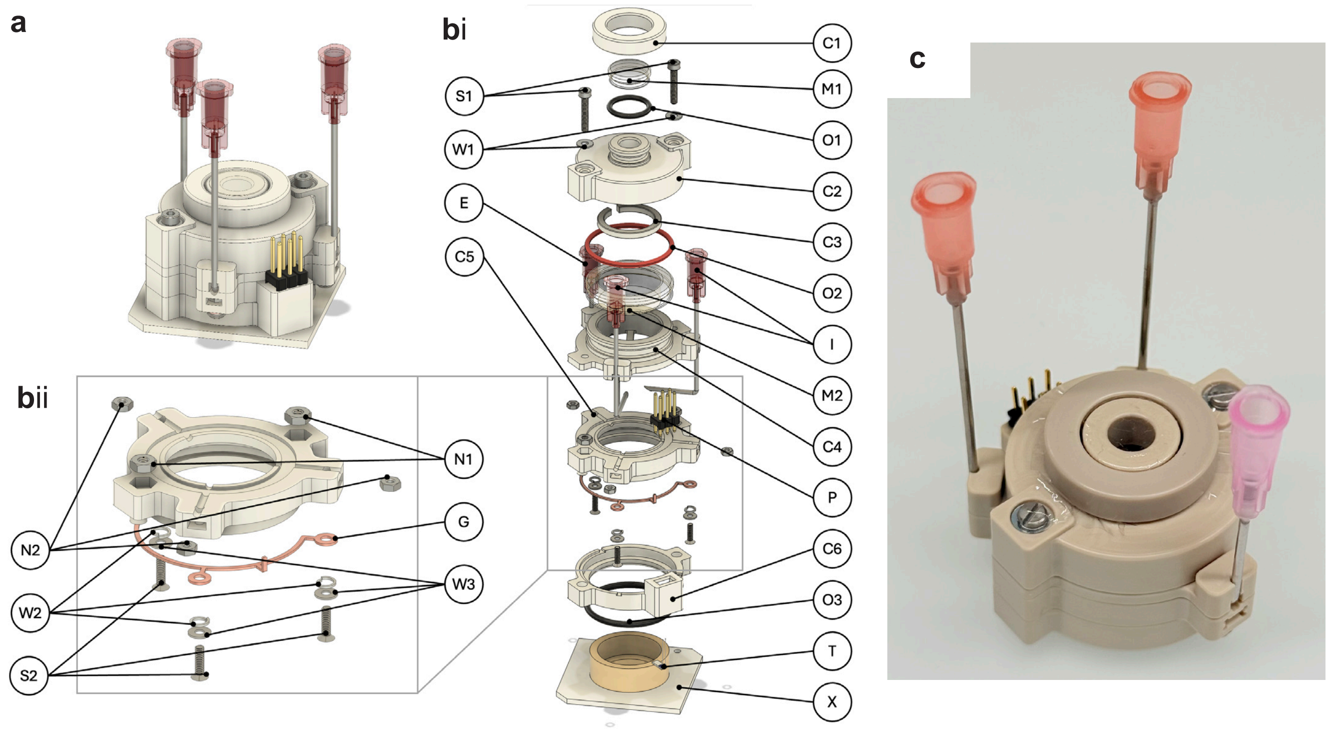


**Fig. S1. CAD design and photo of water compartment lid.** **a**, CAD-rendered overview of the water compartment cap, showing the complete assembly. **b**, (i) Exploded view with (ii) grounding plate details. The lettering corresponds to the label IDs listed in Table 1. **c**, Photograph of the assembled CMOS Cap manufactured from PEEK.

There are 2 membranes (M1, M2 in Fig. S1) positioned above the chip well. The space between the membranes is filled with ultrapure water (Milli-Q, MilliporeSigma, Darmstadt, Germany) and includes an air gap (Fig. S3). To secure the upper membrane, a thin ring (C3 in Fig. S1) is pushed down into the water compartment.

To enable liquid handling within the well while the cap is applied, three stainless-steel needles are integrated into the design: two inlet needles (I in Fig. S1) and one extraction needle (E in Fig. S1). To maximise gas exchange efficiency, the air–liquid interfacial area of the water compartment is designed to be as large as possible. Considering the limited spatial clearance within the inkudock chamber (Fig. S3), the needles are bent upward using a custom bending rig.

To improve needle mechanical stability, the middle component (C4 in Fig. S1) contains snap-fit features that securely hold the needles in two positions. This configuration facilitates assembly and improves robustness during operation. The inlet needles are bent to avoid direct contact with the culture medium, thereby minimising undesired diffusion of compounds through the fluidic system. In contrast, the extraction needle remains in contact with the medium to allow aspiration from the well (Fig. S3).

A grounding plate (G in Fig. S1) is fabricated by laser-cutting a 0.45 mm thick copper sheet into the desired outline. After bending to its final geometry, the plate is integrated into the lower part of the cap and soldered to the connector (P in Fig. S1), as illustrated in Fig. S2. Electrical contact between the grounding plate and the needles is established using screws, washers, and spring washers (S2, W3, W2 in Fig. S1).

For monitoring chip surface temperature, a temperature sensor (T in Fig. S1) is integrated into the cap. The mounting geometry for the sensor within the lower bottom component (C6 in Fig. S1) is specifically designed to optimise its thermal response to surface temperature changes on the CMOS chip. Several geometrical configurations were evaluated, varying the protrusion distance with respect to the bottom surface. The fastest response to temperature fluctuations was achieved when the sensor slightly protruded from the surface. During assembly, the sensor is first bonded into the lower bottom component using adhesive

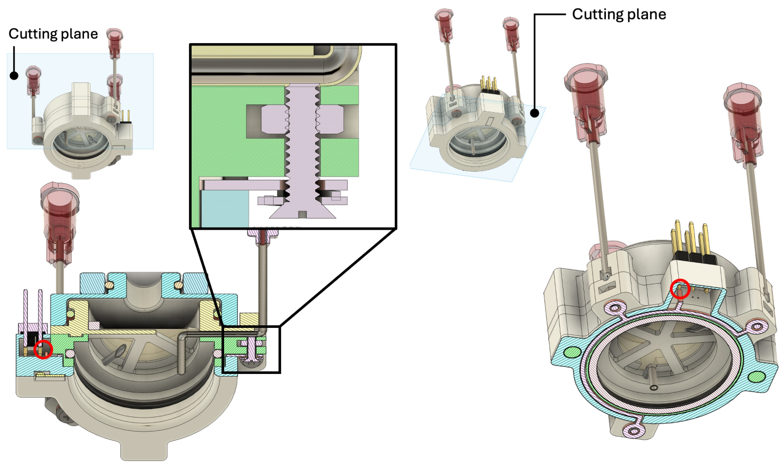


**Fig. S2. CAD renderings of the grounding system shown in two sectional views. The red circles highlight the soldering locations between the grounding plate and the connector.** Left: Vertical section with detail view demonstrating the electrical connection between the needles and the grounding plate established by M1.6 screws. Right: Horizontal section illustrating the position of the grounding plate.

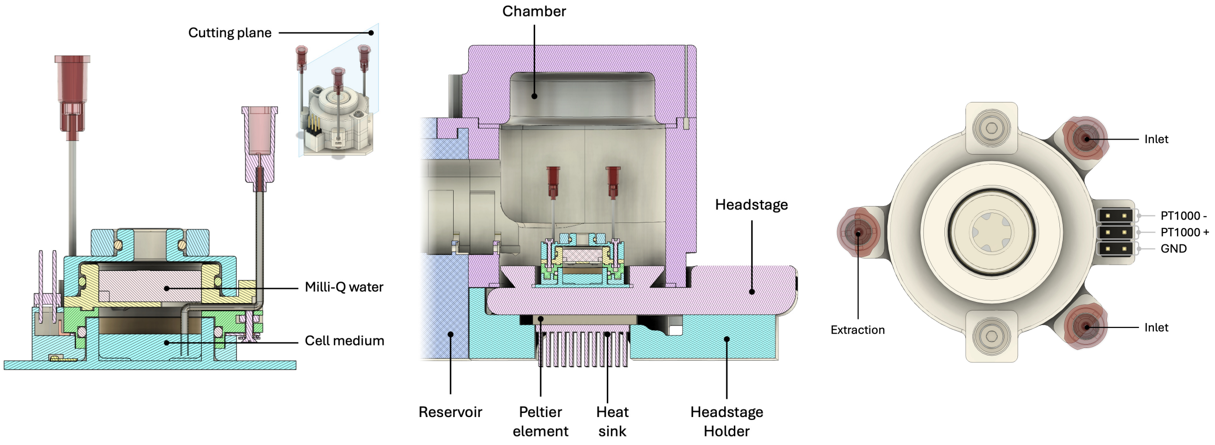


**Fig. S3. Detail views of the water compartment lid and setup.** Left: Sectional CAD view of the water compartment cap mounted on a CMOS chip. Centre: Sectional CAD view of an inkudock chamber containing a CMOS chip with the water compartment cap installed. Right: Top-view CAD rendering of the water compartment cap illustrating the arrangement of liquid-handling and electrical connections.

and subsequently soldered to the connector (P in Fig. S1). The corresponding pin assignments for all components are shown in Fig. S3.
To ensure an airtight seal between the interior of the CMOS chip well and the external environment, a combination of adhesive bonding and O-rings is employed. Adhesive is applied along the glue ridge of the upper-bottom component (C5 in Fig. S1), encapsulating sections of each needle in epoxy and thereby forming a robust, leak-tight interface. The membranes are sealed using o-rings (O1, O2 in Fig. S1). The top ring (C1 in Fig. S1) and top component (C2 in Fig. S1) are compressing the o-rings to achieve a gas-tight seal. Finally, the interface between the CMOS chip well and the water compartment cap is sealed by a press-fit o-ring (O3 in Fig. S1), ensuring complete isolation of the interior of the CMOS chip well.
Stepwise instructions with detailed depictions can be found in the Assembly Instructions.

***A.1.2. Simplified water compartment HD-MEA lid.*** For applications in the standard incubator without recording, for short-term recordings, or in combination with the heated top component (C10 in Fig. S5, see Section A.1.3), the design of the water compartment cap is adapted into a simplified version that omits both the temperature sensor and the liquid-handling needles (Fig. S4). Consequently, a grounding system is no longer required. These simplifications enable a fully mechanical assembly that eliminates all glueing steps, thereby simplifying fabrication and maintenance. While the middle component and the top ring are interchangeable with those of the standard cap, the top component is unique, as the simplified version is also used with standard glass-based MEAs, requiring a reduction in the overall outer dimensions to fit the recording setup.

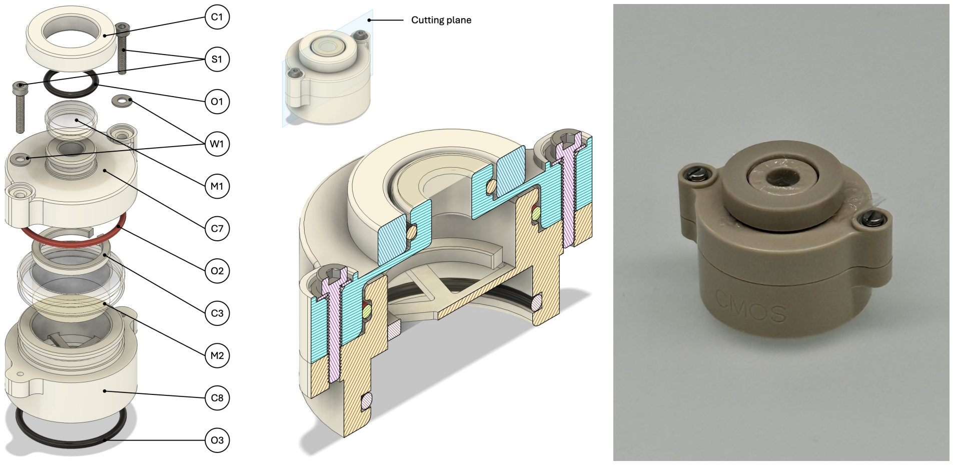


**Fig. S4. CAD design and image of the simplified MEA lid.** Left: Exploded CAD view of the simplified water compartment cap without temperature sensor or liquid-handling needles, showing all major components and o-ring placements. Centre: Section view of the assembled cap illustrating the sealing interfaces and internal geometry. Right: Photograph of the assembled cap manufactured from PEEK. To differentiate it from other cap versions, "CMOS" is engraved laterally.

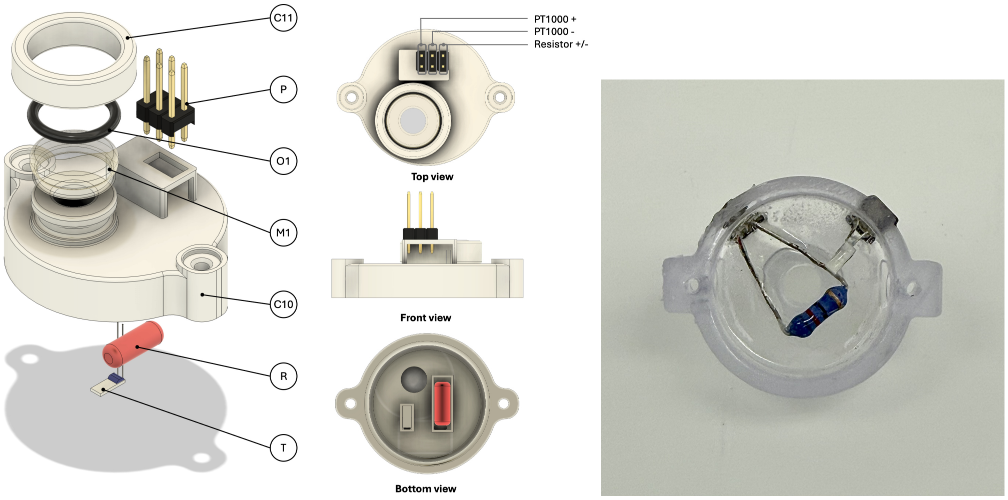


**Fig. S5. CAD design and first prototype image of the heated water compartment lid.** Left: Exploded CAD view of the heated water compartment cap, showing all major components and sealing elements. Centre: Different views of the assembled cap illustrating the internal geometry. Right: Photograph of the assembled top part 3D-printed from resin.

***A.1.3. Heated water compartment HD-MEA lid.*** To replicate the temperature gradient achieved in inkudock, where reservoir heating and recording unit temperature control are combined, within a standard incubator, a heated variant of the top component was developed for use with the simplified water compartment lid (Section A.1.2). A resistor (R in Fig. S5) and temperature sensor (T in Fig. S5) are embedded within a protrusion of the heated top component that extends into the water compartment, enabling active temperature control of the water. In all experiments, the water compartment was maintained at 40 °C while the incubator was set to 35.5 °C. The component was fabricated using resin-based 3D printing (Formlabs, Somerville, MA, USA). The prototype used in experiments (Fig. S5) differs from the final CAD design in the mechanical support and sealing of the embedded components: in the prototype, the resistor and sensor were suspended without structural support and protected only by a thin layer of Araldite 2011, whereas the final design fully encapsulates them for protection of the access wires.

***A.1.4. Water compartment glass-MEA lid.*** The water compartment cap was further adapted for use with standard passive glass-based MEAs. While the MEA well (60MEA500/10iR-Ti-gr, Multi Channel Systems MCS GmbH, Reutlingen, Germany) has the same outer diameter as the MaxOne, its lower height required an adjustment of the bottom component geometry. The middle ring, top, and top ring are compatible with the simplified water compartment cap, ensuring design consistency and simplifying fabrication.

***A.1.5. Leakage test lid.*** The leakage test cap is identical in design to the water compartment cap, except that instead of a middle membrane (M2 in Fig. S1), Middle Ring (C3 in Fig. S1) and a water layer, it contains a solid block of material (C12 in Fig. S6).

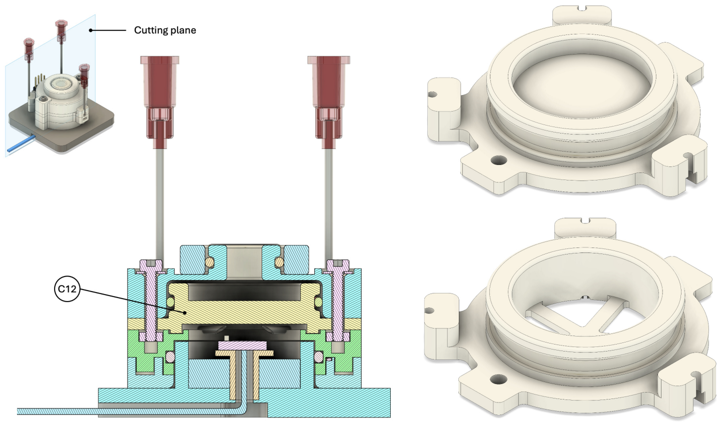


**Fig. S6. CAD design of the cap used for leakage test.** Left: Section view of the CAD model of the leakage test cap. Right: CAD Rendering of the Verification Middle (C12) on the top and the regular Middle Component (C4 in Fig. S1) on the bottom.

### A.2. Custom incubator inkudock.

***A.2.1. Overview.*** The block diagram in Fig. S7shows the different components and their respective boards.

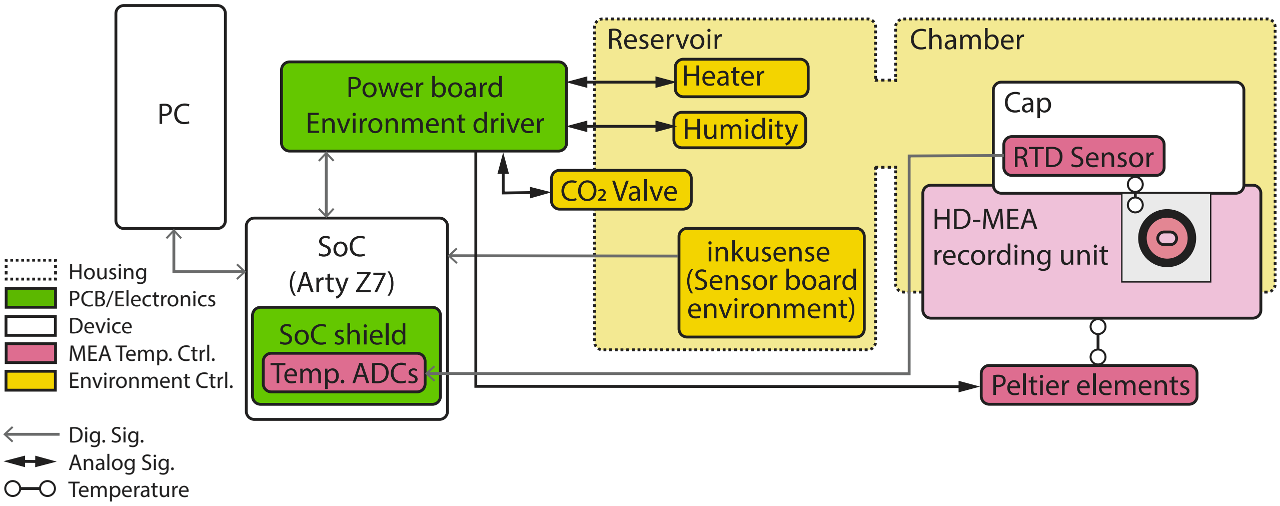


**Fig. S7. Block diagram of inkudock.** The PCBs are adapted from inkube. All active components for environment control are placed inside the reservoir.

***A.2.2. Housing.*** Fig. S8A shows an annotated picture of inkudock with partially mounted chambers.

**Reservoir** The reservoir, depicted in Fig. S8A, carries the sensors and active components for temperature and $CO_2$ control, as well as fans to equilibrate the housing inside. A passive water bath can be used to humidify the environment. The assembly is presented in Fig. S9 and Fig. S10.

**Chamber** The chamber carries no active components. It encloses the HD-MEA and connects to the reservoir, as shown in Fig. S8B.

**Holder** On the holder, the Peltier device is embedded. On top the recording unit is placed. The fan is mounted at the end of the ventilation channel. Details are shown in Fig. S11.

***A.2.3. Electronics.*** The electronics are based on the inkube setup (13). Two custom printed circuit boards (PCBs) are embedded in the setup: the power board and the extended shield. The power board is a revised version of the inkube_pwr. Besides a few minor fixes, the humidity and reservoir heater have been changed to single-ended FET-based output drivers. The extended shield is a combination of the previous inkube_fpga and inkube_bot. It houses the temperature measurement and relays the power board and SoC signals.
Besides the custom PCBs, one replicate of the inkusense board is required for control of the reservoir (47). As heaters, 2 pairs of 2 resistive heaters connected in series are used (RH). For heating and cooling the recording unit, 2 Peltier devices are connected in series (TD) and attached to a heat sink (HS) with thermally conductive foil (HF). Further, 2 fans per mounted recording unit help distribute the air inside the reservoir and chamber and dissipate heat from the heat sink (F).

### A.3. Validation sensing setups.

***A.3.1. Sensor dummy chip.*** To evaluate the $CO_2$ diffusion characteristics of the water compartment cap, a custom test setup (hereafter referred to as the *test well*) was fabricated. The test well replicates the geometry of the CMOS chip and includes a central platform that holds a sensor board capable of recording relative humidity, temperature, and $CO_2$ concentration. The sensor board was developed in a previous project where it was used to control an incubator setup (47). The sensor board cables are routed through a hole in the platform and well and sealed with Araldite 2011 adhesive to ensure an airtight connection (Fig. S12).

***A.3.2. Sensor ring.*** The sensor ring incorporates the same PT1000 embedding design as the water compartment lid, but features a reduced structure around the well, forming a thin ring that allows a commercial single-membrane cap to be placed on top (Fig. S13).

**A.4. Component and Supplier Details.** Tables 1 and 2 summarise the components used in the water compartment cap and the inkudock incubator, respectively. PCB design files, 3D CAD models, and detailed component-level BOMs are available in the ETH Research Collection (see data and material availability statement).

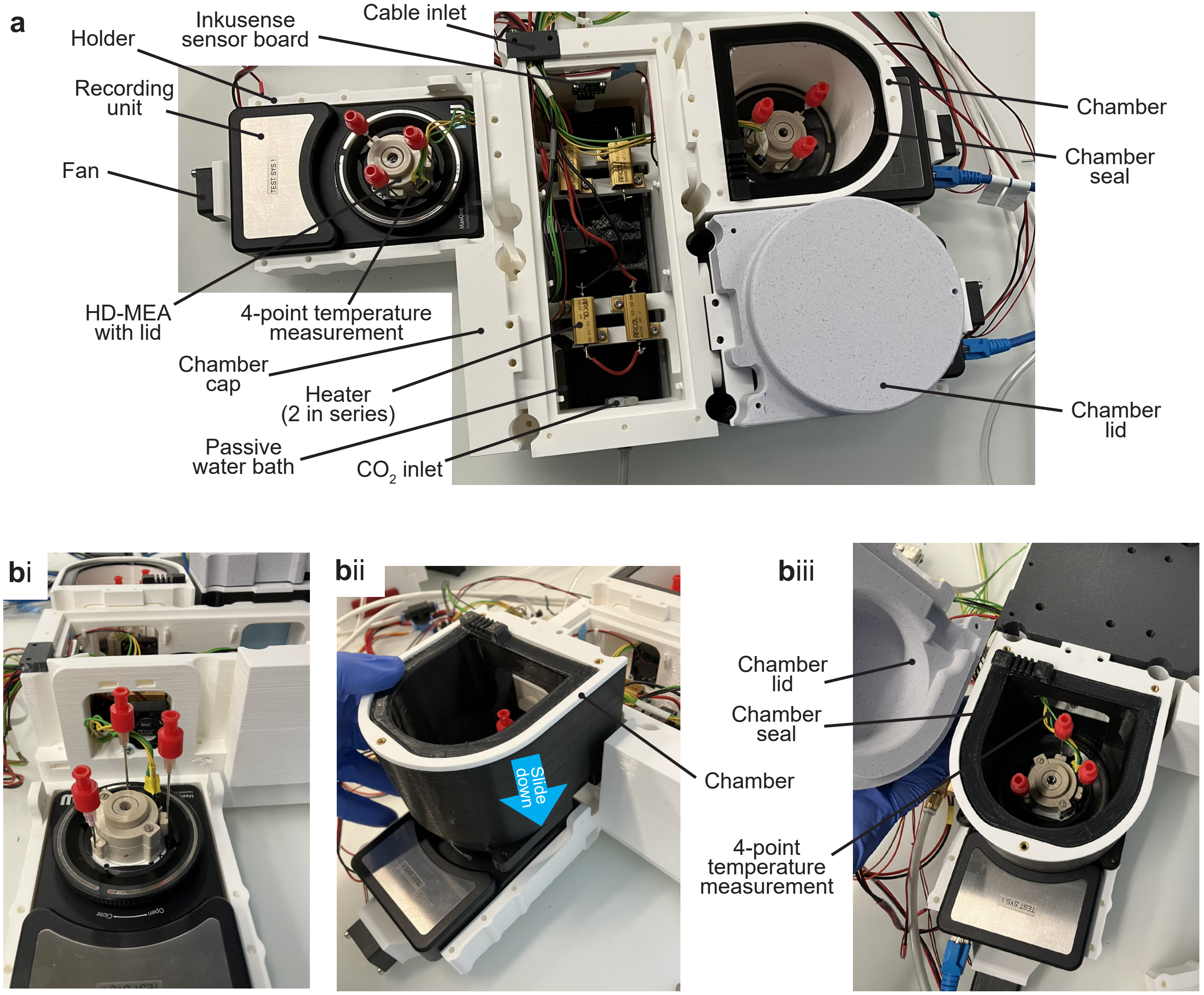


**Fig. S8. inkudock overview a**, Top view of the assembled inkudock with one closed chamber on the bottom right, a chamber with the lid removed on the top right, a recording unit without the chamber on the top left, and a closed connection port on the bottom left. **b**, Chamber mounting. **(i)** Recording unit with mounted chip aligned to a port. **(ii)** The chamber can be mounted by sliding in from the top to hold the recording unit in place and connect the chamber interior to the reservoir. **(iii)** Finally, the chamber seal is placed, and the lid can be attached.

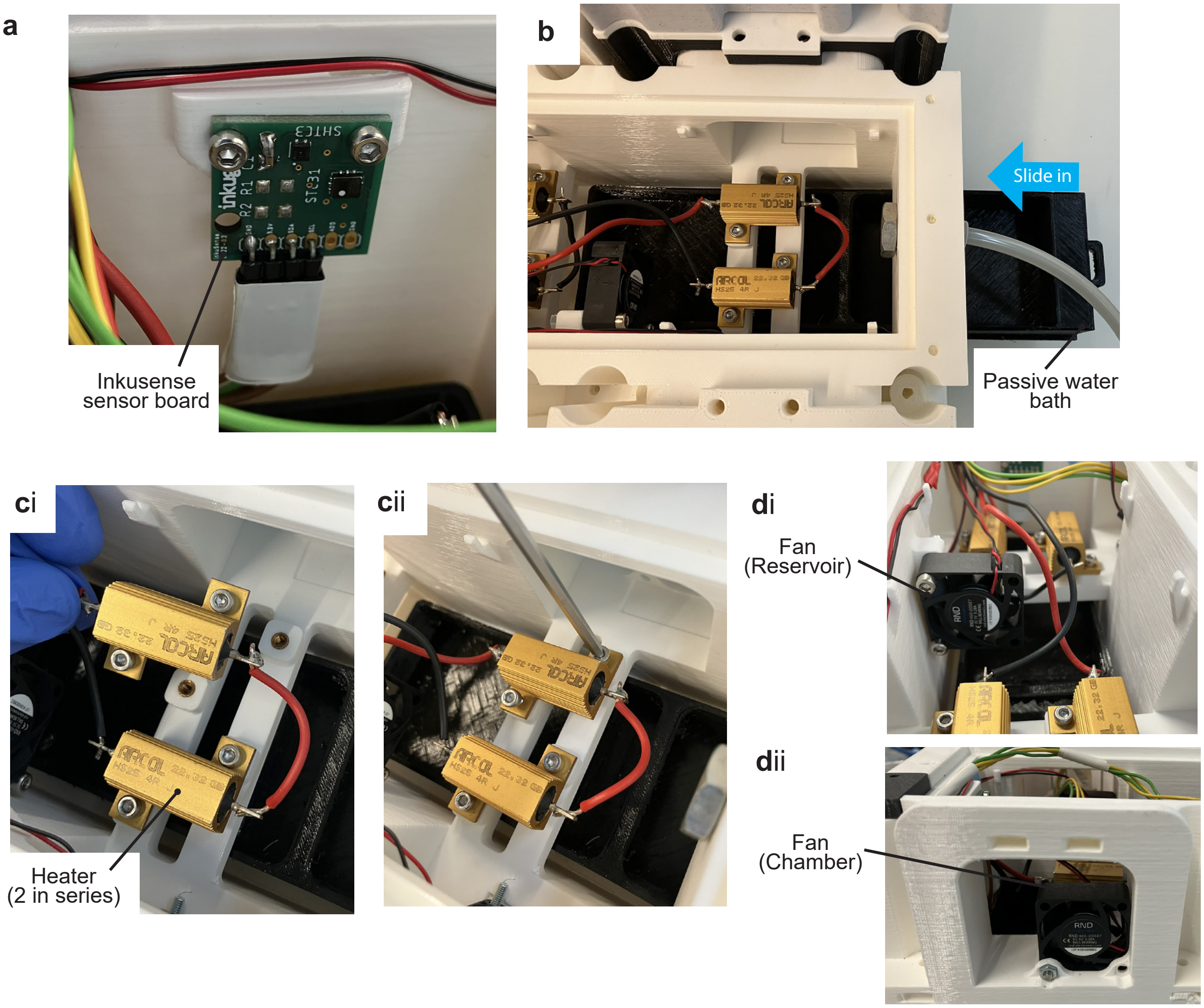


**Fig. S9. Mounting of components in the reservoir.** **a**, The inkusene board is attached with 2 M2.5x6 screws with counter hex nuts. **(i)** The water bath is slid in from the side. **c**, Mounting the resistive heaters. **(i)** For heating of the reservoir, four 4 Ω resistive heaters (RH in 2) are placed, split into 2 branches of 2 resistors in series. **(ii)** Two M3 thread inserts per resistor are placed in the holes and the resistors are mounted using 2 M3x8 screws. **d**, Fan mounting. **(i)** One central fan (F in 2) is placed in the reservoir, with the airflow pointing towards the $CO_2$ inlet, by using a M3x18 screw and a counter hex nut. **(ii)** Per chamber, one fan (F in 2) is used to balance temperature gradients and attached to the inkudock port with one M3x20 screw.

| ID | Name | Specification | Manufacturer / Supplier |
|---|---|---|---|
| *Cap Parts (CNC machined)* | | | |
| C1 | Top Ring | PEEK / PMMA | In-House |
| C2 | Top | PEEK / PMMA | In-House |
| C3 | Middle Ring | PEEK / PMMA | In-House |
| C4 | Middle | PEEK / PMMA | In-House |
| C5 | Upper Bottom | PEEK / PMMA | In-House |
| C6 | Lower Bottom | PEEK / PMMA | In-House |
| C7 | Top Glass MEA | PEEK / PMMA | In-House |
| C8 | Bottom CMOS NS/NT | PEEK / PMMA | In-House |
| C9 | Bottom Glass MEA | PEEK / PMMA | In-House |
| C12 | Verification Middle | PMMA | In-House |
| *Cap Parts (3D printed)* | | | |
| C10 | Heated Top | BiomedClear Resin | In-House |
| C11 | Heated Top Ring | BiomedClear Resin | In-House |
| *Seals and Membranes* | | | |
| O1 | Top O-Ring | Viton 75 Shore, $10\times1.5$ mm (0103-004528) | Kubo, CH |
| O2 | Middle O-Ring | Viton 75 Shore, $23\times1.5$ mm (0103-212057) | Kubo, CH |
| O3 | Bottom O-Ring | MVQ 70 Shore, $24\times2$ mm (0102-194567) | Kubo, CH |
| M1 | Small Membrane | ALA MEA-MEM-SHEET, 50 Gauge | MCS GmbH, DE |
| M2 | Large Membrane | ALA MEA-MEM-SHEET, 50 Gauge | MCS GmbH, DE |
| *Fasteners* | | | |
| S1 | M2×12 Screw | Pan Head, Slotted, V4A SS | toolcraft AG, DE |
| S2 | M1.6×6 Screw | Pan Head, Slotted, A2 SS | RS Components, CH |
| N1 | M2 Nut | SS, DIN 934 | RS Components, CH |
| N2 | M1.6 Nut | A2-304 SS, 3.20 mm, DIN 934 | RS Components, CH |
| W1 | M2 Washer | Stainless Steel | RS Components, CH |
| W2 | M2 Spring Washer | Zn-Plated Steel, $0.5\times2.2\times4.4$ mm | RS Components, CH |
| W3 | M1.6 Washer | Stainless Steel | RS Components, CH |
| *Other Components* | | | |
| P | Connector | 2×3 Pin Header, 2.54 mm pitch | RS Components, CH |
| I | Inlet Needle | Sterican, $1.2\times50$ mm, sharp | B. Braun, DE |
| E | Extraction Needle | Sterican, $1.2\times40$ mm, blunt | B. Braun, DE |
| T | Temp. Sensor | PT1000, NB-PTCO-170 | TE Connectivity / DigiKey |
| G | Grounding Plate | Copper Sheet, $300\times300\times0.45$ mm | In-House, laser cut |
| *MEA Systems* | | | |
| X1 | CMOS HD-MEA | MaxOne+ HD-MEA chip | MaxWell Biosystems, CH |
| X2 | Recording system | MaxOne recording system | MaxWell Biosystems, CH |

**Table 1.** Bill of materials for the water compartment cap.

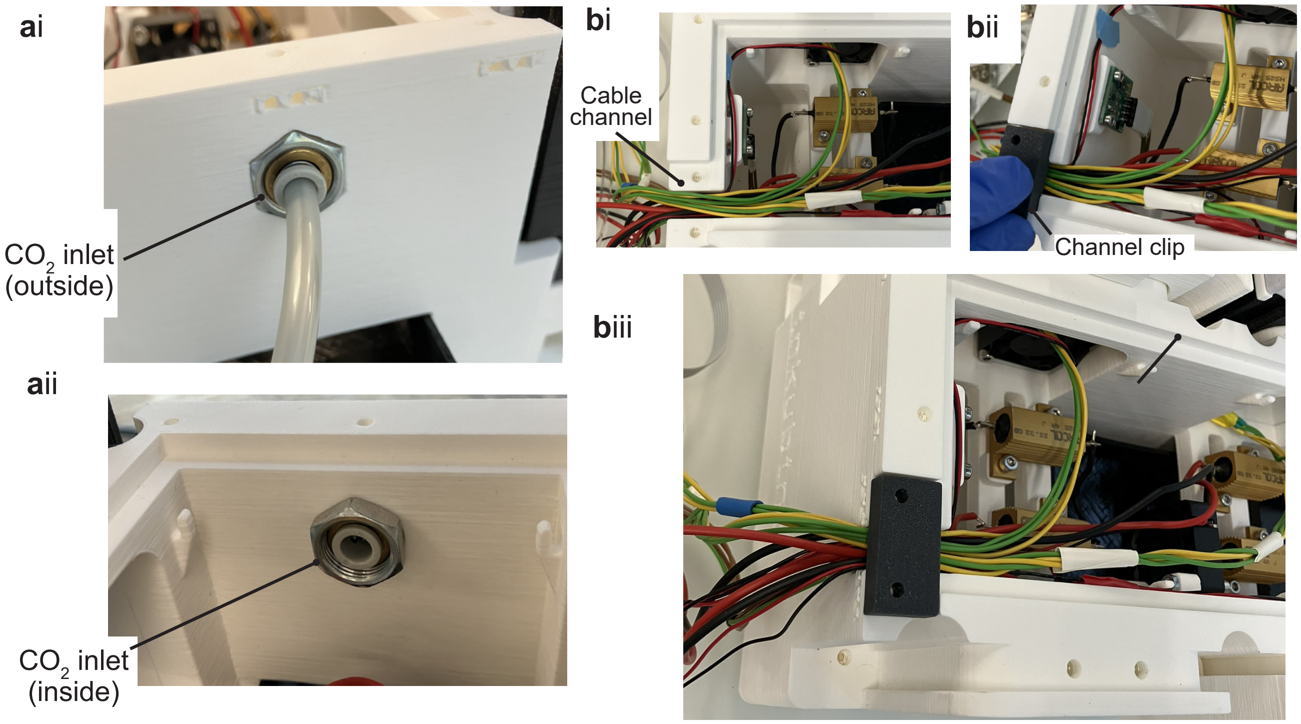


**Fig. S10.** Mounting inlets and outlets of the inkudock. **a**, $CO_2$ connection. **(i)** The transduction thread is mounted from outside into the hole. The tube supplying the $CO_2$ from the pulsing valve can be plugged in. **(ii)** The thread is secured from the inside with a counter hex nut. **b**, All cables leading inside the inkudock are aligned to the cable channel next to the sensor board. **(ii)** and **(iii)** A clip is used to secure the cables.

| ID | Name | Specification | Manufacturer / Supplier |
|---|---|---|---|
| *Reservoir (3D printed, PLA)* | | | |
| CAD1 | Reservoir housing | PLA, 3D printed | In-House |
| CAD2 | Reservoir lid | PLA, 3D printed | In-House |
| CAD3 | Water bath | PLA, 3D printed | In-House |
| CAD4 | Reservoir seal | PLA, 3D printed | In-House |
| CAD5 | Port cap | PLA, 3D printed, seals unused chamber ports | In-House |
| *CMOS MEA Chamber and Recording Holder (3D printed, PLA)* | | | |
| CAD6 | Recording holder | PLA, 3D printed | In-House |
| CAD7 | Chamber | PLA, 3D printed | In-House |
| CAD8 | Insulation chamber | TPU, 3D printed | In-House |
| CAD9 | Chamber lid | PLA, 3D printed | In-House |
| *Printed Circuit Boards* | | | |
| PCB1 | Power board | 4-layer, power_board v4.4 | Custom, see Research Collection |
| PCB2 | Extended shield | 2-layer, extended_shield v4.4 | Custom, see Research Collection |
| PCB3 | Inkusense board | 2-layer, inkusense v4.2 | Custom, see (47) |
| *Electronic Components* | | | |
| SoC | FPGA/SoC board | Arty Z7-20, Zynq-7000 Z7020 (410-346-20) | Digilent / DigiKey |
| PSU | Power supply | DT80PW090D, 9 V | TDK-Lambda / DigiKey |
| *Thermal Management (per holder)* | | | |
| TD | Peltier device | TEC1-07901, 50×20 mm (×2, series) | Wellen Technology Co., China |
| HS | Heat sink | ICK S 45×45×20, pin type | Fischer Elektronik, DE |
| HF | Thermal foil | WLFT 404, 40×40 mm | Fischer Elektronik, DE |
| F | Fan | RND 460-00087, 30×30×10 mm, 5 V (×2) | Distrelec AG, CH |
| *Environment Control* | | | |
| RH | Resistive heater | HS25 4R J (×4, 2×2 series) | ARCOL / Distrelec |
| V | $CO_2$ valve | PVQ13-6M-06-M5-A | SMC Pneumatics / Distrelec |

**Table 2.** Bill of materials for the inkudock incubator.

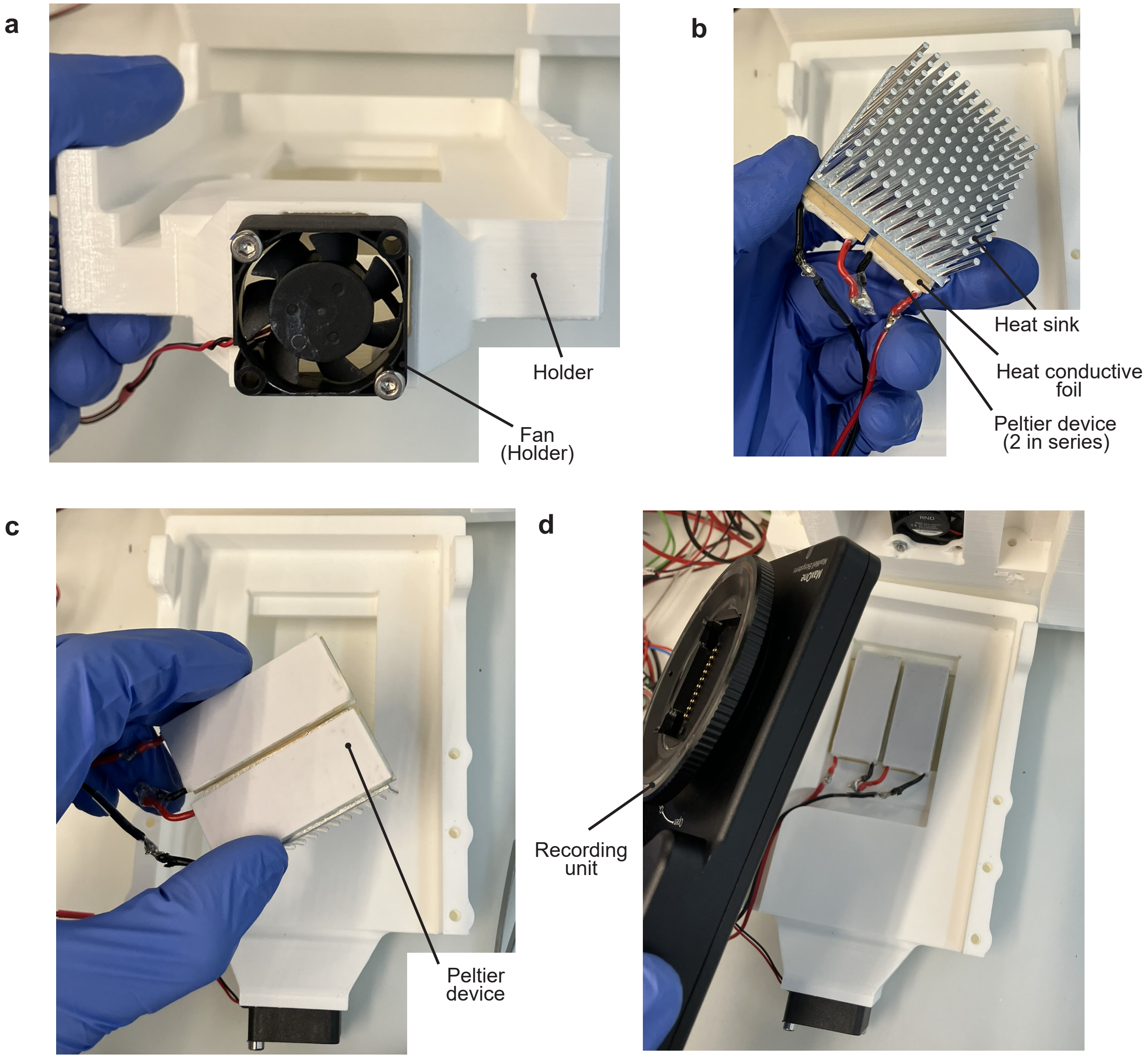


**Fig. S11. Assembly of the holder.** **a**, The fan (F in 2) is mounted at the end of the ventilation channel, with 2 M3x15 screws and M3 thread inserts, pointing its airflow towards the Peltier device. **b**, Peltier device are mounted with a thermally conductive adhesive foil on top of a heat sink, with the labelled side of the Peltier devices exposed towards the top. For sufficient power, 2 Peltier devices are connected in series (be aware of the polarity). **c**, The assembly is then placed in the gap of the holder and the cables are lead out. **d**, Finally, the MaxOne recording unit is placed on top of the holder.

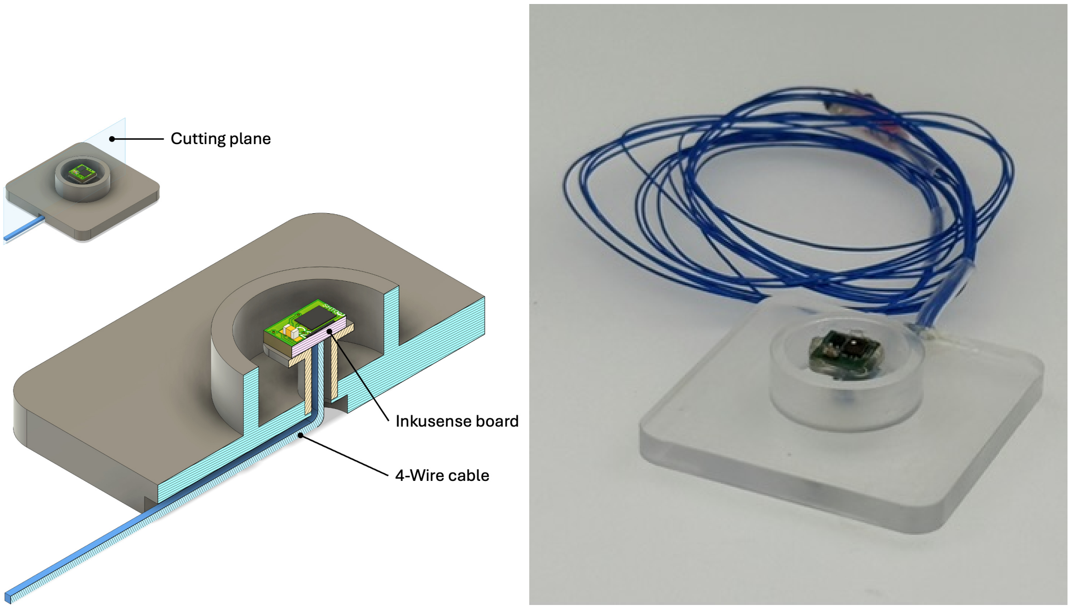


**Fig. S12.** Left: Section view of the CAD model of the test well illustrating the well geometry and chamber configuration. Right: Photograph of the test well fabricated from PMMA.

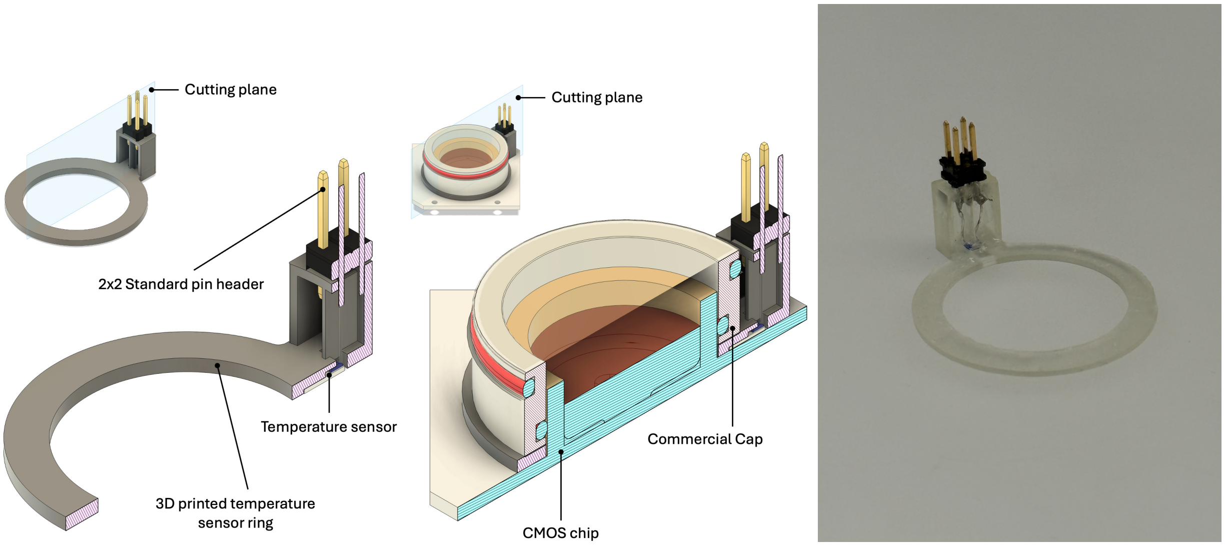


**Fig. S13.** Left: Section CAD view of the sensor ring. Centre: Section CAD view of the sensor ring mounted on a CMOS chip with a single-membrane lid mounted on top of it. Right: Photograph of the sensor ring.

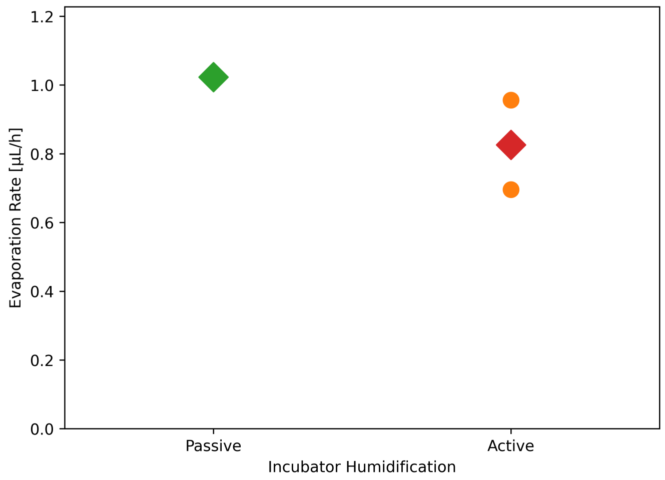


**Fig. S14. State-of-the-art evaporation in incubator.** Evaporation was measured over 48 h from 3 HD-MEAs in commercial incubators. One was placed in an incubator with passive humidification, and two were placed in incubators with active humidification. For the latter, the individual measurements are presented as dots and the mean is marked with a diamond. All MEAs were placed with 1 mL of water and a commercial membrane lid was mounted. MEAs were weighed before and after 48 h to determine the volume of evaporated liquid.

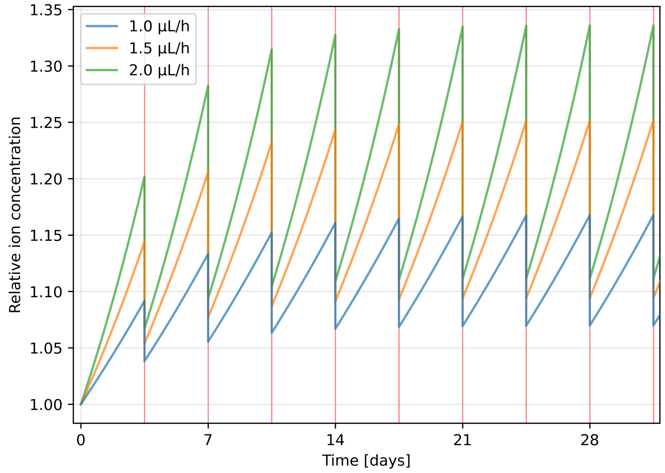


**Fig. S15. Simulated relative ion concentration in the culture medium over time under a half-medium exchange protocol.** Each cycle consists of 3.5 days of evaporation followed by removal of half the well volume (500 µL from a 1000 µL starting volume) and replenishment with fresh medium to restore the original volume. Curves are shown for three constant evaporation rates (1.0, 1.5, and 2.0 µL/h). Medium exchange is indicated with red vertical lines.

### Supplementary Note B: Additional experimental data

**B.1. Evaporation in incubator with membrane lid.** Fig. S14 shows evaporation from HD-MEAs in a Petri dish and with a commercial membrane lid in commercial incubators to provide a reference for expected evaporation from HD-MEAs following state of the art protocols.

**B.2. Ion concentration change due to half-medium exchange.** To estimate the cumulative effect of evaporation on extracellular ion concentrations, we modelled the medium as a discrete recurrence over exchange cycles of 3.5 days. Within each cycle, evaporation removes only water at a constant rate, thereby concentrating all solutes by a factor $V_0/(V_0 - V_{\mathrm{evap}})$, where $V_0 = 1000\,\mu\mathrm{L}$ is the starting volume and $V_{\mathrm{evap}}$ is the total volume lost to evaporation per cycle. At the end of each cycle, 500 µL of the remaining medium are removed, and the starting volume $V_0$ is restored by adding fresh medium (500µL + evaporated volume). For the common evaporation rates in an incubator with a membrane lid (1.0–2.0 µL/h), steady-state concentrations are elevated by at least approximately 7–11 % (right after medium exchange) relative to fresh medium after 30 days.

**B.3. Temperature calibration.** Measurements during calibration of the temperature control loop showed a chamber temperature of approximately 43 °C when the reservoir was set to 45 °C (Fig. S16C). To achieve physiological conditions for the cells, the relation between medium and HD-MEA surface temperature was determined. The sensor in the culture medium was therefore controlling the Peltier device with a set temperature of 35.5 °C, which was established to correspond to 38.5 °C on the HD-MEA surface. The fact that the set temperature is higher than the medium temperature is presumably explained by the sensor placement. As the sensor is mounted on the lid and pressed onto a corner of the MEA chip, it is still strongly influenced

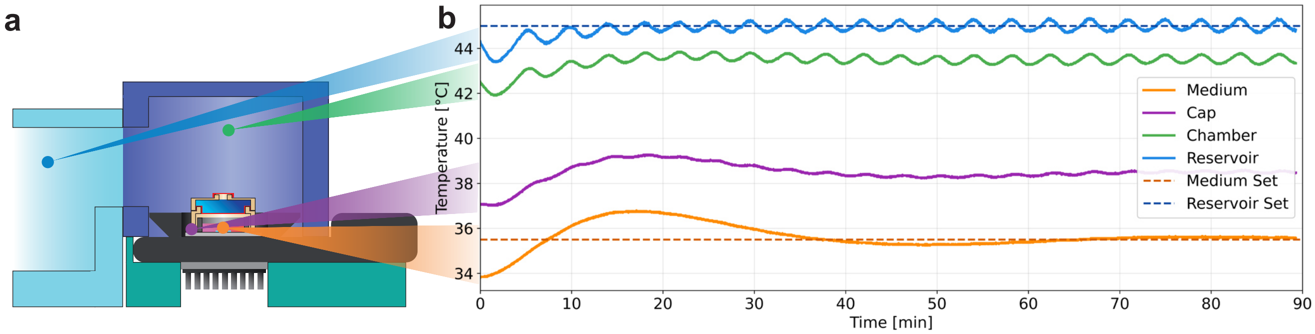


**Fig. S16.** Temperature control calibration of the incubation setup. (i) Measurement locations of the temperature sensors. An independent loop controls the reservoir temperature and is set to 45 ℃ (blue). A second sensor measures the temperature inside the chamber (green). In order to determine the set temperature for the second control loop with the temperature sensor on the chip (purple), a sensor is placed inside the liquid with a set temperature of 35.5 ℃ (orange). In steady state, the chip temperature is 38.5 ℃, which is further used as the set temperature for the chip sensor (purple).

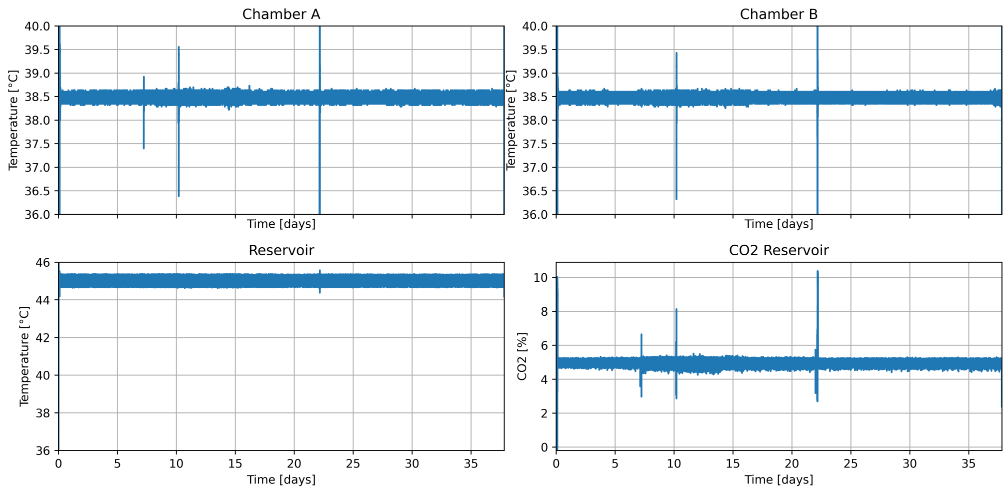


**Fig. S17.** Culturing conditions during the long-term recording. The temperature in the chambers is set to 38.5 °C, with interrupts in Chamber A during opening of the chamber lid on day 7, the water compartment refill on day 10 and the medium exchange on day 22. The reservoir temperature is unaffected and stable at 45 °C. The $CO_2$ concentration is set to 5 % with small deviations at chamber opening and the 10% settings for 30 min at start and after medium exchange.

by the temperature of the chamber environment. The recording unit, in contrast, is presumably colder as the bulk of the material is outside of the chamber, and the lid serves as a good insulator. This would then lead to a temperature gradient across the HD-MEA, with a lower temperature in the centre and a higher temperature outside. For all further experiments, the HD-MEA surface was controlled to this value, and the temperature sensor in the liquid was omitted to reduce the risk of contamination.

**B.4. Environmental data during long-term recording.** Temperature settings and $CO_2$ concentration during the long-term recording in inkudock are shown in Fig. S17. The medium exchange after 22 days is visible as a fluctuation in the temperature and the $CO_2$ is increased to 10 % for 30 min after restart. After 10 days, a refill of the water compartment is performed, leading to a dip and small overshoot in the chip temperature.

**B.5. Medium comparison.** Culture medium was sampled from MEA cultures with the water compartment cap after 14 days without medium exchange and from cultures after 3 days in the incubator with a membrane cap. Both cultures were seeded at a density of 150 k cells.

Glucose concentration was measured using a commercial blood testing glucometer (CONTOUR NEXT 20438293, Ascensia Diabetes Care, Parsippany, NJ, USA). Medium was diluted 1:1 with PBS to fit within the glucometer's measurement range. Linear regression through all data points with a fixed intercept at 100% (fresh medium) yielded a glucose consumption rate of -1.36 %/day ($R^2$ = 0.679). After 14 days, glucose levels remained at approximately 80 % of the initial concentration. Given the high glucose concentration in Neurobasal medium (25 mM), glucose depletion is unlikely to affect culture viability.

The NanoDrop One Microvolume UV-Vis Spectrophotometer (ThermoFisher) was used to characterise the protein content in the different medium conditions. The measurements are shown in Fig. S19 and Fig. S20

Protein electrophoresis was performed using the NuPAGE™ Bis-Tris gel system (NP034B, Invitrogen) with the same media, and gels were stained with SimplyBlue™ SafeStain (LC6060, Invitrogen). Protein size was estimated using the PageRuler™ Prestained Protein Ladder, 10 to 180 kDa (26616, Thermofisher) as calibration. Protein loads of 10 and 20 µg were analysed in

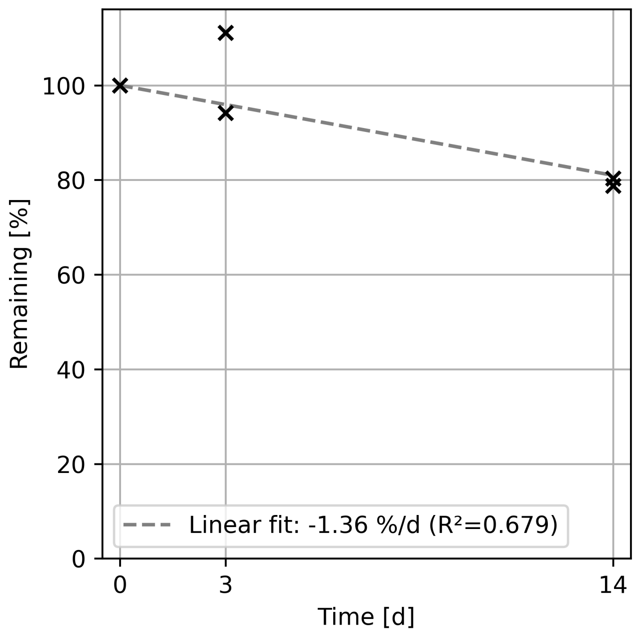


**Fig. S18.** Glucose concentration remaining in culture medium over time. Fresh medium (day 0) serves as 100% reference. Individual measurements from day 3 and day 14 are shown with crosses. Dashed line represents linear fit forced through 100% at day 0.

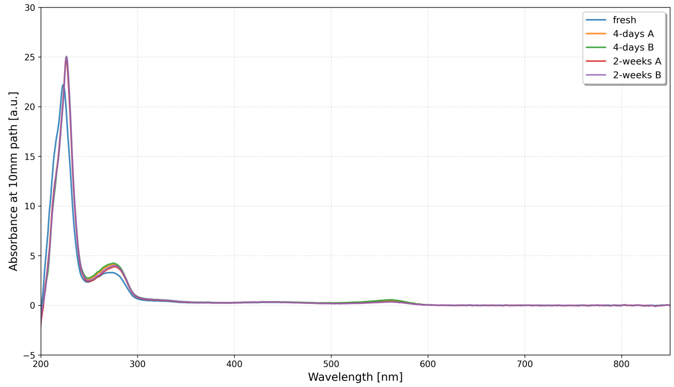


**Fig. S19.** Nanodrop analysis of samples.

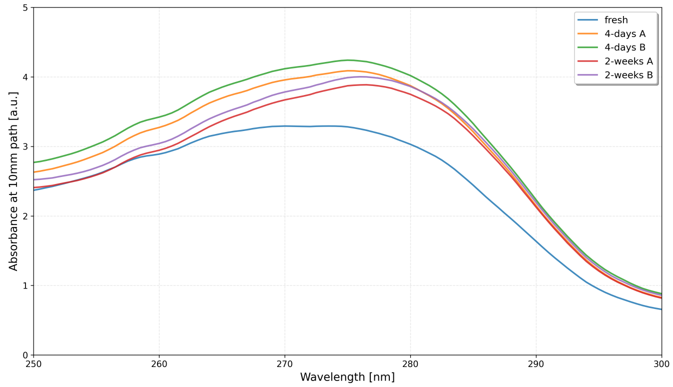


**Fig. S20.** Zoom in on the nanodrop analysis, small changes at the 275 nm peak are visible.

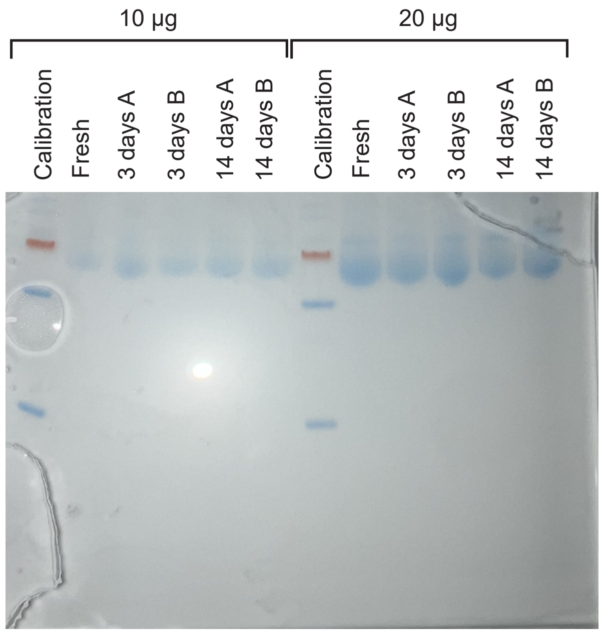


**Fig. S21.** Protein Electrophoresis (from left to right): calibration, fresh medium, 2 samples after 3 days, 2 samples after 14 days at 10 $\mu$g of protein and the same samples at 20 $\mu$g. No apparent differences in protein content are visible.

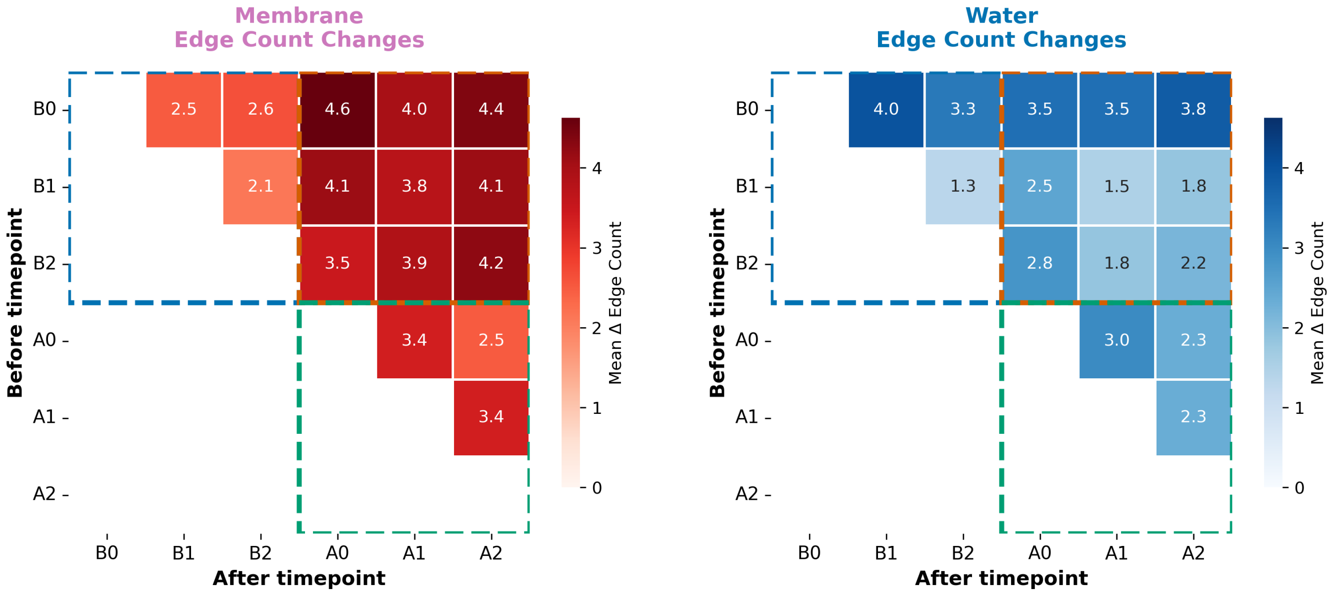


**Fig. S22.** Edge count changes across all timepoint pairs. Mean edge count changes (sum of absolute differences per source-target connection, normalised TE $\geq 1 \times 10^{-4}$) for all transitions. Networks with weakly connected component $\geq$3 nodes included. Boxes highlight within-period (blue, green) versus across-exchange (orange) transitions for membrane and water conditions.

Fig. S21.

**B.6. Medium exchange impact TE graph.** The full data for the plot in Fig. 2is shown in Fig. S22.

**B.7. Culture moved between conditions.** A cell culture with a microstructure of 12 networks, each with 2 seeding wells connected via 10 parallel microchannels, was recorded starting at 49 DIV. The mean firing rate per network is shown in Fig. S23. The culture was initially recorded with the water cap in inkudock following a full medium exchange. At day 4 and day 12, half of the culture medium was exchanged. After the second half exchange (day 12), the culture was moved to a standard incubator without recording. At day 15, a brief recording was taken, followed by a full medium exchange. The culture was then recorded for 2 h in the inkudock. The water cap was then exchanged with a commercial membrane cap, and it was transferred to the incubator, where recording continued until firing ceased. Notably, firing rates remained remarkably stable during the inkudock recordings (days 0–12 and day 15), with gradual increases over the first 12 days. In contrast, upon transfer to the

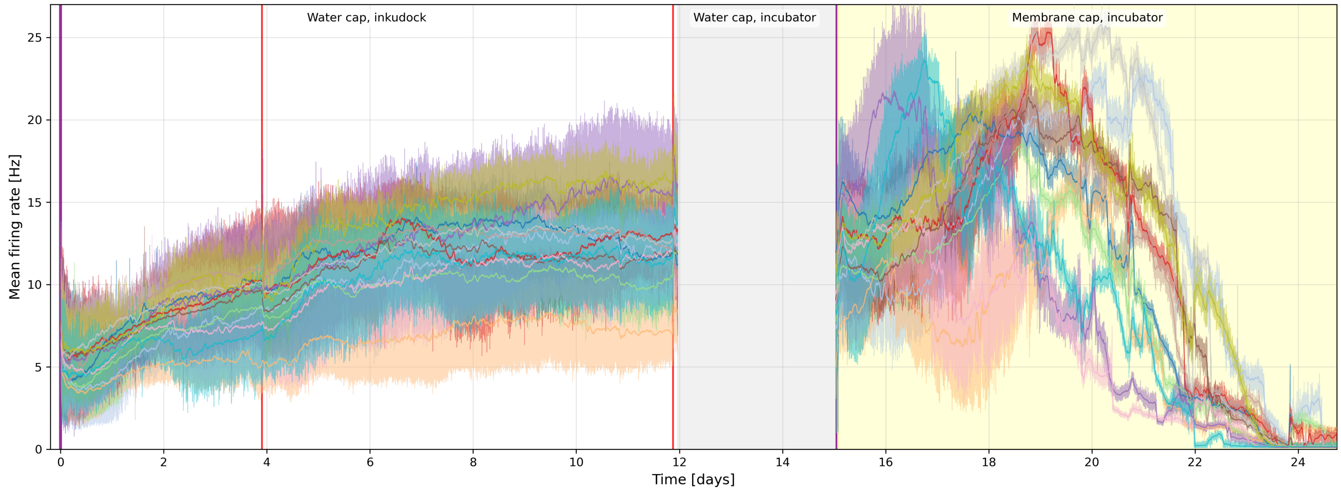


**Fig. S23.** Firing rate of a cell culture during medium exchange and different recording conditions. Electrodes were selected in the microchannels. Activity is processed by network, binned into 10 s and a 3-point median is applied (faint lines) to avoid a drop due to overlap of bins with the data saving breaks. The thin lines show an average across 60 min, excluding the bins during data saving.

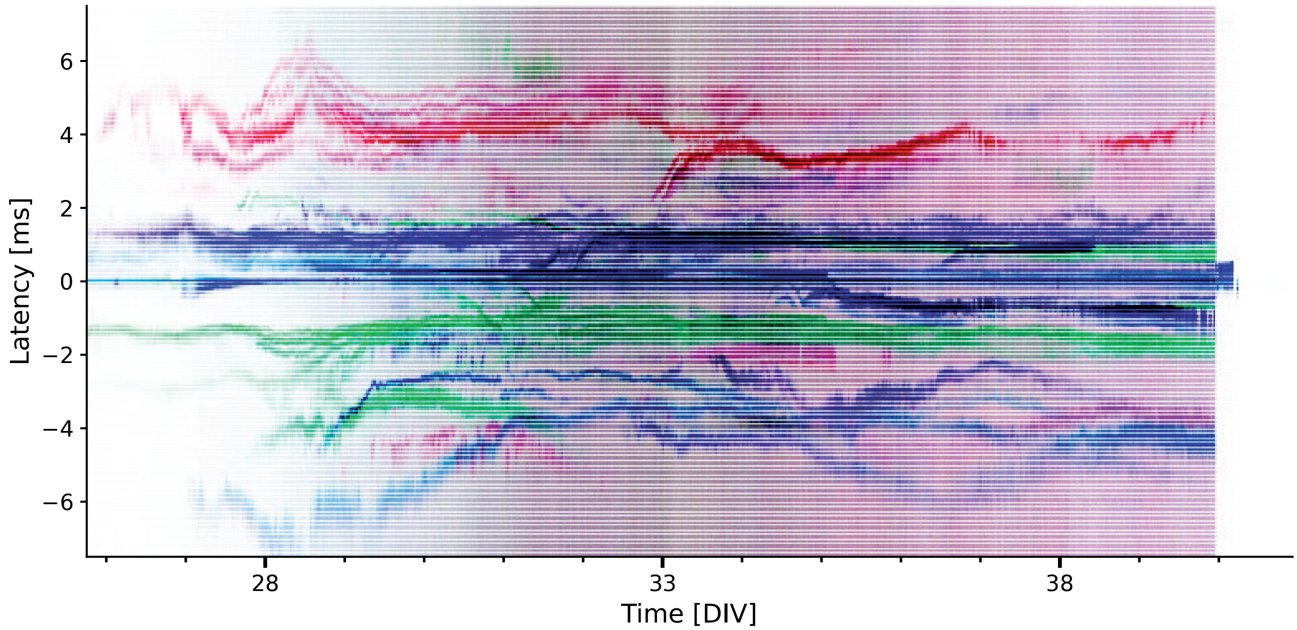


**Fig. S24.** Additional MEA recorded for 14 days without medium exchange.

standard incubator (day 15 onwards), all networks exhibited rapid perturbations within hours followed by an increase in firing rate between 17-19 days and a decline afterwards.

**B.8. Additional network STTRPs.** The HD-MEA was prepared as described in Methods. On 25 DIV, a full medium exchange was performed, the water compartment lid was mounted, and a continuous recording was started.

**B.9. Full clustering.** Patterns sampled from the patterns identified through clustering are shown in Fig. S26.

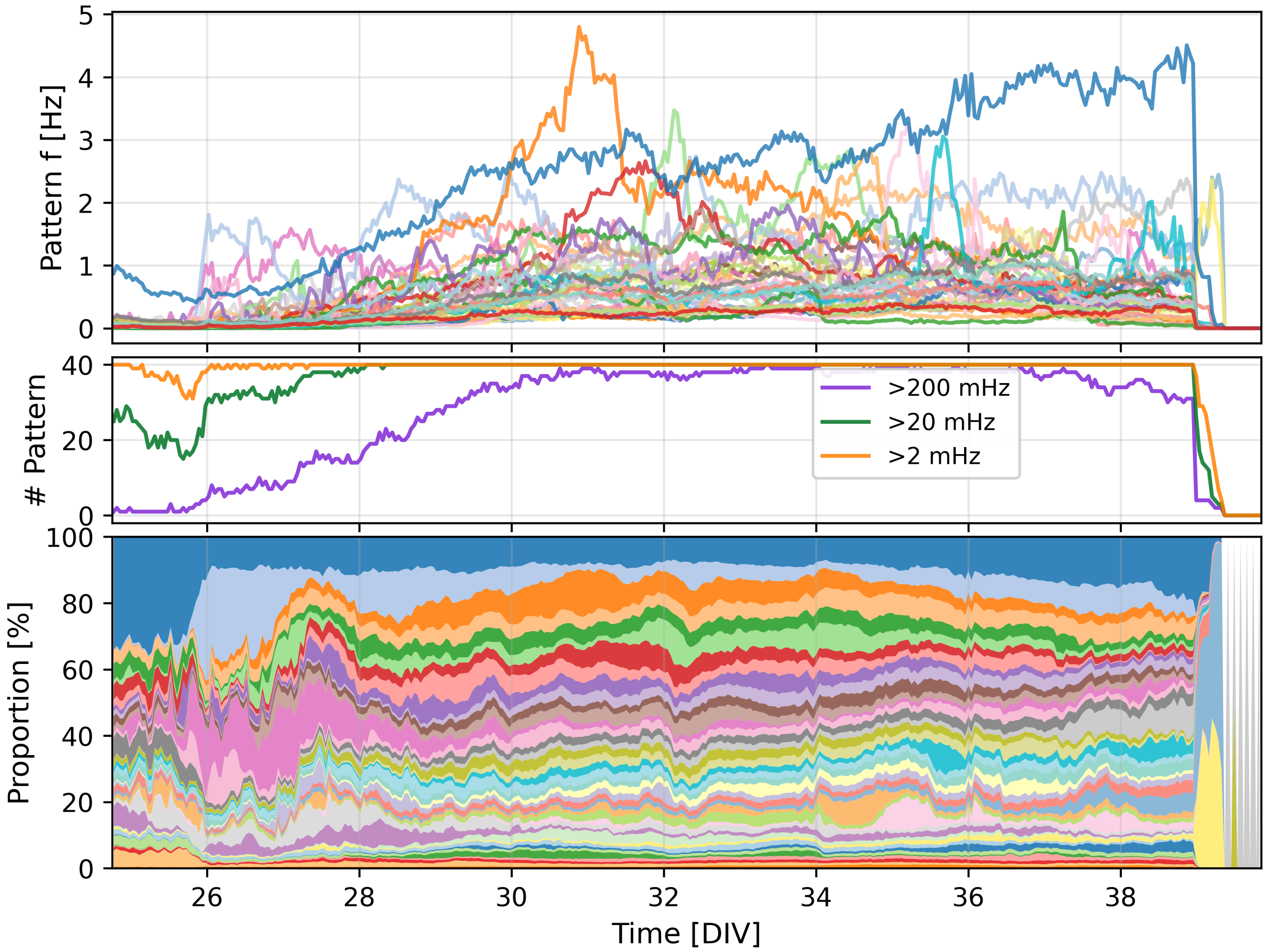


**Fig. S25.** Patterns detected on the network from Fig. S24 analysed as in Fig. 4e-g.

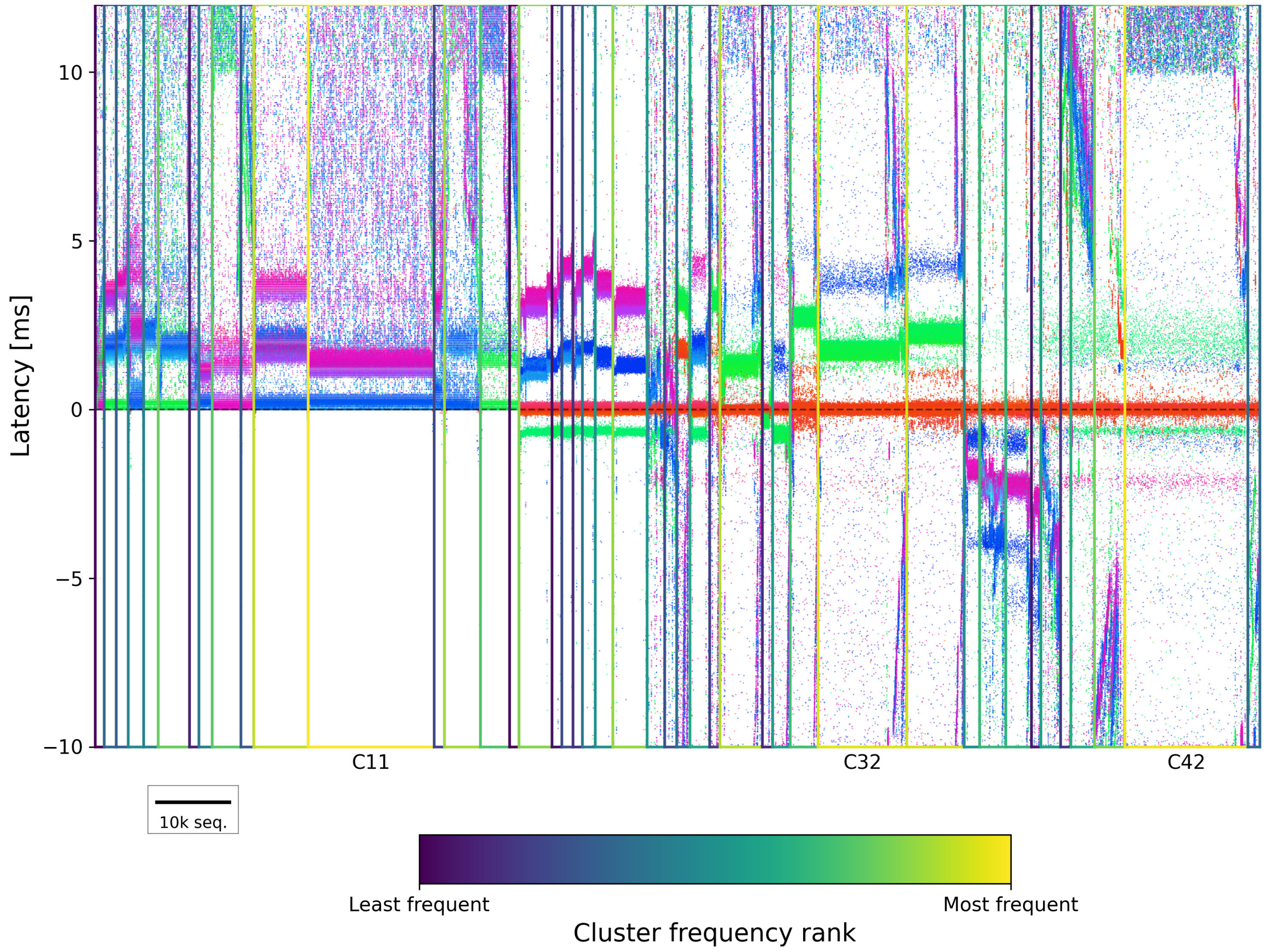


**Fig. S26.** Raster plot of 200,000 randomly sampled spatiotemporal spike sequences, sorted by the first principal component within each of the 44 clusters (labelled with C). Cluster rectangles are coloured by frequency rank (viridis: bright = most frequent). Scale bar: 10,000 sequences.
